# Ultraconserved elements and machine learning classifiers enable robust phylogenetics and taxonomy in model and non-model nematodes

**DOI:** 10.1101/2025.05.06.652396

**Authors:** Laura Villegas, Lucy Jimenez, Jöelle van der Sprong, Oleksandr Holovachov, Ann-Marie Waldvogel, Philipp H. Schiffer

## Abstract

Nematodes are among the most diverse animals, yet only around 28,000 of an estimated one million species have been morphologically described. Their small size, morphological simplicity, and cryptic diversity complicate phylogenetic analyses. Traditional morphological and single-locus molecular approaches often lack resolution for both recent and ancient divergences. To address these limitations, we developed the first ultraconserved elements (UCEs) probe sets for two nematode families: Panagrolaimidae, a group of non-model organisms with limited genomic resources when compared to model taxa, and Rhabditidae, which includes the model species *Caenorhabditis elegans*. Our probe sets targeted 1,612 loci for Panagrolaimidae and 100,397 for Rhabditidae. In vitro testing recovered up to 1,457 loci in Panagrolaimidae, supporting robust phylogenetic reconstruction. Results were largely consistent with previous analyses, except for one strain reclassified as *Neocephalobus halophilus* BSS8. Using machine learning, we determined the minimum number of loci needed for accurate genus-level classification. For Rhabditidae, XGBoost achieved high accuracy with just 46 loci. For Panagrolaimidae, 39 loci were most informative. Our UCE-based approach offers a scalable and cost-effective framework for phylogenomics, enhancing taxonomic resolution and evolutionary inference in nematodes. It is well suited for biodiversity assessments and shallow, field-based sequencing, expanding research possibilities across this ecologically important phylum.

## 4 Introduction

Nematodes (Phylum Nematoda) are one of the most diverse and abundant animal taxa, occupying nearly all ecosystems on earth and playing crucial ecological roles, such as regulating microbial and fungal communities, facilitating nutrient cycling, and contributing to soil health and decomposition processes as well as being parasites of plants and animals, including humans (M. Blaxter, 2011; van den Hoogen et al., 2020). Nematode taxonomic diversity remains poorly characterized, with recent estimates of the total number of species in the phylum ranging between half a million and up to 10 million species (Hodda, 2022). As only a little over 28000 species are formally described, an immense ‘taxonomic gap’ not only limits our understanding of biodiversity, but also constrains our ability to explore the evolutionary history and diversification of this group. Better resolution of nematode phylogeny is furthermore crucial for advancing the understanding of ecological dynamics, e.g., in soils and sediments, where nematode diversity can be linked to patterns on the environmental scale (Villegas et al., 2024).

Resolving phylogenetic placement of species is the basis for reconstructing evolutionary relationships, shedding light on lineage diversification, biogeography, and adaptation. Current approaches, such as morphology-based classification, are often limited by cryptic diversity, convergent evolution, and the need for specialized taxonomic expertise. Similarly, traditional molecular markers (e.g., 18S rRNA, 28S rRNA, COI and others) may lack sufficient phylogenetic resolution to reliably distinguish closely related species, while some nematode taxa exhibiting particularly high molecular evolution rates, which can in turn produce skewed or incongruent topologies in phylogenetic reconstructions (M. L. Blaxter et al., 1998; Van Megen et al., 2009). This issue highlights the importance of developing new methods for phylogenetically robust species placement, with a focus on the careful selection of informative loci. Several sequencing methods have emerged as a complement or alternative to amplicon sequencing, like whole genome sequencing (WGS), transcriptome sequencing, target sequence capture, and restriction-site associated sequencing (RAD-seq). Each of these methods offers distinct advantages. WGS enables the comprehensive investigation of complex genomic traits and adaptive processes (Lu et al., 2025), allowing the study of coding and non-coding regions and the analysis of structural variation. However, this comes at increased costs, and data analysis can be computationally intensive. RAD-seq provides a lower cost alternative for population and species-level studies when compared to whole genome approaches by focusing on a reduced subset of the genome (K. R. Andrews, Good, Miller, Luikart, & Hohenlohe, 2016); however, determining orthology relationships remains challenging. Missing data due to mutations at restriction sites, and potential non-independence of adjacent loci due to linkage disequilibrium are some of the potential drawbacks of this approach (Rubin, Ree, & Moreau, 2012).

For targeted sequence capture methods, specific genomic regions are enriched for sequencing. In this way, a large set of orthologous loci can be selected with probes, making the method particularly useful for phylogenetics and evolutionary studies (M. R. Jones & Good, 2015). However, a major drawback is the requirement of prior genomic or transcriptomic data availability for probe design. Consequently, successful implementation depends on the availability of high-quality genomes, which may be limited for non-model organisms. One common family of bait sets targets highly conserved genomic regions, among them ultra-conserved elements (UCEs), which recover sets of loci that are highly conserved and thus can be captured across divergent groups of organisms. The presence of highly conserved regions (cores) that are anchored between highly variable informative sites (flanking regions) makes it a cost-efficient approach that captures both deep phylogenetic relationships and recent divergences, enabling high-resolution phylogenomic reconstructions (Faircloth et al., 2012). UCEs’ approaches have been successfully implemented in a wide range of lineages (Blaimer et al., 2015; Erickson, Pentico, Quattrini, & McFadden, 2020; Gilbert et al., 2015; Quattrini et al., 2017; van der Sprong et al., 2023; Winker, Glenn, & Faircloth, 2018). Beyond evolutionary studies, UCE-based analyses can also provide critical insights into ecologically relevant patterns, such as population structure, connectivity, and demography, which are essential for conservation planning and bio-monitoring efforts (Duckett, Calder, Sullivan, Tank, & Carstens, 2023).

Ultimately, the most suitable method depends on the focus taxonomic group and specific research objectives, as different approaches vary in their effectiveness depending on genome complexity, evolutionary scale, and data requirements. In this study, we designed two ultraconserved elements (UCEs) bait sets for nematodes, particularly focusing on the Rhabiditidae (to which the model nematode *C. elegans* belongs to) and Panagrolaimidae (non-model species) families. Using Rhabditidae for method benchmarking, we obtained phylogenetic reconstructions based on UCE loci that were congruent with those previously proposed in the literature based on orthologs from genome-wide analysis. For the Panagrolaimidae family, for which only a few genomic resources are available compared to Rhabditidae, we show that using UCEs and target capture provides a high-resolution phylogenetic reconstruction. Additionally, we developed a machine learning framework to identify the most informative UCEs for the taxonomic assignment of genera, offering a scalable and efficient tool for future ecological and evolutionary studies.

The genomic era has generated high-dimensional datasets that pose significant challenges to traditional taxonomy, particularly in groups with high diversity, like nematodes (M. Blaxter, 2011; Hodda, 2022). To effectively interpret these complex data, it is essential to adopt computational approaches that can uncover intricate patterns within genomic variation. The application of machine learning (ML) techniques offers a promising way to analyse UCE presence-absence and highlights the ability of ML algorithms to leverage and identify informative features and build robust predictive models (Libbrecht & Noble, 2015). Notably, presence–absence data have also proven effective for phylogenetic inference, recovering meaningful evolutionary signal even in complex datasets (Natsidis, Kapli, Schiffer, & Telford, 2021). The necessity to select relevant features within these complex datasets requires efficient feature selection methods, which allow for improved model interpretability and reduced dimensionality without compromising the accuracy (Guyon & Elisseeff, 2003; Saeys, Inza, & Larrañaga, 2007). Therefore, the integration of ML and UCE analysis represents a promising strategy to address taxonomic challenges in nematodes, offering a scalable and objective framework for genus-level classification (Lv, Li, Wang, & Zou, 2023). The use of presence–absence patterns of UCEs in this context is of particular interest, as these binary patterns may carry important phylogenetic signal that could improve genus-level classification and contribute to a more accurate understanding of evolutionary relationships. This approach is especially valuable in groups where traditional taxonomic expertise is limited or insufficient, providing a new way to explore complex phylogenetic structures within nematodes.

## 5 Materials and methods

### 5.1 Phylogenomic analysis of the family Panagrolaimidae: in-silico and in-vivo testing

#### 5.1.1 Bait set design and in-silico testing

##### Bait set design

Phyluce 1.7.2 was used for the bait set design, the “Tutorial IV: Identifying UCE Loci and Designing Baits To Target Them” was followed (phyluce.readthedocs) (Faircloth, 2015; Faircloth et al., 2012). Eight base genomes were tested for the bait set design: *Panagrolaimus* sp. PS1159 (GenBank accession number: GCA 901765195.1), *Panagrolaimus* sp. ES5, *Panagrolaimus* sp. PS1579 (GenBank accession number: GCA 901779485.1), *Panagrolaimus kolymaensis* (GenBank accession number: GCA 028622995.1), *Panagrolaimus* sp. LJ2400 (GenBank accession number: GCA 024447215.1), *Panagrolaimus* sp. LJ2406 (GenBank accession number: GCA 024447205.1), *Panagrolaimus* sp. LJ2414 (GenBank accession number: GCA 024447195.1) and *Panagrolaimus superbus* (GenBank accession number: GCA 901766145.1).

Bait sets were individually designed using each of the base genomes and further assessed based on the quality of the base genome, the number of conserved loci consistently found between exemplar taxa, and the final probe count (for further synthesis).

Illumina reads for each of the strains used as base genomes were obtained through the Sequence Read Archive (SRA), with the exception of *Panagrolaimus kolymaensis*, which was de novo sequenced using a NovaSeq Illumina platform 2×150 bp by Illumina UK Company Limited. Publicly available re-sequencing data for the other strains tested were obtained through the Sequence Read Archive (SRA) (supplementary table 3).

Reads were mapped against the base genomes using stampy (v 1.0.32) (Lunter & Goodson, 2010), allowing for sequence divergence ≤ 5. Unmapped reads were removed using samtools (v 1.18) (Danecek et al., 2021). Filtered BAM files were converted to bed format, and overlapping intervals were merged using bedtools (v 2.31.1) (Quinlan & Hall, 2010) for putatively conserved regions. Repetitive intervals and ambiguous bases were removed using phyluce probe strip masked loci from set, intervals shorter than 80 bp and 25% of the regions were masked using the same tool. Sequences with the identified UCEs were extracted using phyluce probe get genome sequences from bed specifying a buffer region of 160 bp. Temporary bait sets targeting those sequences were obtained using phyluce probe get tiled probes with a tilling density set to three, using the −masking, –remove−gc flags, potentially problematic baits with over 25% repeat content and GC content outside of 30-70% were also removed. Duplicate baits were screened and removed using phyluce probe easy lastz and phyluce probe remove duplicate hits from probes using lastz.

The baits were then aligned against each of the tested base genomes using phyluce probe run multiple lastzs sqlite with an identity value of 50%. Baits that matched multiple contigs were removed. Using phyluce probe slice sequence from genomes, fasta sequences from each exemplar taxa were extracted, buffering each locus to 180 bp. The final bait design was obtained using phyluce probe get tiled probe from multiple inputs. The baits were designed with a tilling-density of three, baits with 25% masking were removed, and two probes targeting each locus were designed. Duplicate baits were removed (with an identity value of 50%) using phyluce probe easy lastz and phyluce probe remove duplicate hits from probes using lastz.

The selection of the preferred bait set was based on three criteria: the quality of the base genome, the total number of loci targeted, and the total number of probes (see “ProbeDesignPS1159 selection” in the Zenodo repository – 10.5281/zenodo.15395838). The different bait lists were further examined by the bioinformatics team from the company BIOCAT to confirm the number of baits (sequences), GC content, and size of the probes. The selected bait set was synthesized using myBaits® Custom (1–20,000) target capture kit by Daicel Arbor BioSciences (Ann Arbor, MI, USA).

#### 5.1.2 Sampling, DNA extraction and sequencing

Nematodes were harvested from cultures in agar plates by washing off plates with 2mL of nuclease-free water, pelleted by centrifuging at 4 degrees for 5 min at 3000g. Nematodes were kept as laboratory cultures at 15°C on low-nutrient agar plates inoculated with OP50 *(E. coli)*. Reproductive mode assessment has previously been performed by (Lewis et al., 2009; Villegas et al., 2024) on a set of the strains here tested. DNA extractions for in-vitro testing of the baits were performed using the Quick-DNA Microprep Plus Kit (catalogue number: D4074) manufacturer’s protocol for tissue samples. The protocol was modified by increasing the proteinase K volume to 20*µ*L per sample and incubating overnight for 20 hours. One entire plate was used per strain for DNA extraction. After DNA extraction for 21 strains (supplementary table 5), library preparation and sequencing were done at the Cologne Genome Center for Genomics (supplementary table 6). DNA extractions, library preparation, and sequencing were performed following the Arbor+ TruSeq Nano DNA protocol (version 02.2024). For DNA fragmentation, 600 ng of high-quality DNA per sample was processed using a Bioruptor (Diagenode) with a target fragment size of 350 bp (range: 310–350 bp), and quality was assessed using a 4200 TapeStation (Agilent) with D1000 ScreenTape. Library preparation was performed using the TruSeq Nano DNA Library Prep Kit (Illumina) with 100 ng of fragmented DNA and eight PCR cycles. Library quality was checked with a 4200 TapeStation (Agilent) D1000 ScreenTape. Pooled libraries were prepared with a total of 2 µg DNA per pool, corresponding to either seven samples at approximately 223 ng each or nine samples at approximately 286 ng each, and volume was reduced to 7 µL by evaporation. Hybridization was performed for 24 hours at 62°C, followed by capture wash steps at 62°C and elution in 25 µL of Buffer E. Post-capture PCR was performed using reagents from the TruSeq Nano DNA Kit, using 10 PCR cycles following the Arbor-PCR program (initial denaturation at 95°C for 3 minutes, followed by 10 cycles of 98°C for 20 seconds, 60°C for 30 seconds, 72°C for 45 seconds, and a final extension at 75°C for 5 minutes). Final libraries were cleaned using a 1:1 ratio of AMPure XP beads and eluted in 30 µL of EB buffer, with quality control performed using a 4200 TapeStation (Agilent). Sequencing was carried out on a NovaSeq 6000 Illumina sequencer (S4 flowcell) with 2×151 bp paired-end reads.

#### 5.1.3 Analysis of sequencing data: in-vivo testing and phylogenomic reconstruction

Sequencing data resulting from target capture was analysed following the “Tutorial I: UCE Phylogenomics” (phyluce.readthedocs.io) with Phyluce 1.7.3. Sequencing adapters were trimmed using Cutadapt (v 4.9)(M. Martin, 2011). Quality of the trimmed reads was assessed using FastQC (v0.11.2) (S. Andrews, 2010). Assemblies for each strain were conducted using Spades (v 3.14.1) (Prjibelski, Antipov, Meleshko, Lapidus, & Korobeynikov, 2020) incorporated in phyluce (phyluce assembly assemblo spades). Quality of the assemblies was checked using phyluce assembly get fasta lengths. Genomes available through NCBI for the Panagrolaimidae family were used to obtain a more comprehensive phylogenetic reconstruction of this family and test the robustness of using UCEs as a smaller subset of the genome for phylogenetic reconstructions (supplementary table 4). UCE loci from genomes were harvested following the “Tutorial III: Harvesting UCE Loci From Genome”(phyluce.readthedocs. Genomes were converted into 2bit format using faToTwoBit (v 445 and 472) (R. M. Kuhn, Haussler, & Kent, 2012). The UCE probe set was aligned to the genomes, and the fasta sequences corresponding to the loci were extracted using phyluce probe run multiple lastzs sqlite and phyluce probe slice sequence from genomes. The extracted FASTA sequences were then processed along with the UCE loci obtained from the Spades assemblies for the sequenced target capture data.

For known triploid taxa, we used the program dedupe.sh from the BBMap suite (Bushnell, 2014). This tool compares all sequences to each other and removes those that are highly similar, keeping only the longest version of each sequence. We set the minimum identity threshold to 90% (minidentity=90), meaning that any sequences sharing 90% or more nucleotide identity were considered duplicates and collapsed into a single representative. This step was used to avoid including multiple copies of the same UCE locus that may result from allelic variation, assembly errors, or the presence of homologous copies in triploid genomes. We then matched the assembled contigs and available genomes of each strain to the UCE baits using phyluce assembly match contigs to probes. The UCE loci were then extracted using phyluce assembly get fastas from match counts. In the case of *Panagrolaimus kolymaensis*, the contigs were fragmented into 2000 bp long and the assembly file was split into 3 different files using a round-robin method since harvesting UCEs from this genome yields a high number of duplicated UCEs due to the triploid nature, and the deduping method was not useful.

The UCE loci, both harvested from genomes and from target capture data, were aligned and edge-trimmed using phyluce align seqcap align. The alignments were then cleaned using phyluce align remove locus namefrom files. Subsequently, a data matrix of 60% occupancy was created using phyluce align get only loci with min taxa (each UCE that was present in at least 60% of the taxa analyzed). The alignments were concatenated using phyluce align concatenate alignments. The resulting alignment was used as input for IQ-tree (v 2.3.6) (Minh et al., 2020), a maximum likelihood inference was made using ultra-fast bootstrap (1000 bootstrap replicates) with the Model Finder Plus feature for automatic selection of the best substitution model for the dataset. The resulting phylogenetic tree was visualized using TreeViewer (v 2.2.0) (Bianchini & Śanchez-Baracaldo, 2024) using the *Further transformation* module to root the tree and change rename labels from accession number and sequencing code to strain or species name, and the *Plot actions* module to display the legend with bootstrap values and add the scale bar.

#### 5.1.4 Species reassessment based on morphological and molecular evidence

For the strain previously identified as *Panagrolaimus detritophagus* BSS8, morphometric measurements were taken and compared with those of closely related species to accurately determine its taxonomic identity. Measures included, among others, body length, vulva location, lip region width, stoma length, corpus length, pharyngeal region length, and tail length in both females and males.

For light microscopy, specimens were relaxed using heat, fixed in a cold 4% formaldehyde solution, gradually transferred to pure glycerine using a slow evaporation method, and mounted on permanent slides in glycerine, with paraffin wax used to support the coverslip. Nine male and nine female individuals isolated from current cultures were compared to closely related species: *Neocephalobus halophilus* Paetzold, 1958 (Paetzold, 1958), *Neocephalobus aberrans* Steiner, 1929 (Steiner, 1929a) and *Panagrolaimus orthomici* Korentchenko, 1992 (Korentchenko, 1992) as well as to the description of the original wild population of the same “*Panagrolaimus* population II” (Bostrom, 1988), see supplementary files.

### 5.2 Ultraconserved elements for accurate species delimitation: test case using ***Caenorhabditis***

#### Bait-set design and in-silico testing

Phyluce 1.7.3 was used for the bait set design, the “Tutorial IV: Identifying following UCE and Designing Baits To Target Them” was followed from the official documentation phyluce.readthedocs. Only one base genome was tested in this case, this corresponded to *Caenorhabditis elegans* (Bristol N2) (GenBank accession number: GCA 000002985 3). The probe design was performed in the same way as for Panagrolaimidae, with the exception of using artificially generated reads in this case, using illumina art as specified in the phyluce tutorials instead of real sequencing data. For the generation of this family’s bait set, the following *Caenorhabditis* species were used to identify putatively conserved regions: *Caenorhabditis brenneri* (GenBank accession number: GCA 964036135.1), *Caenorhabditis briggsae*(GenBank accession number: GCA 021491975.1), *Caenorhabditis elegans* (Bristol N2) (GenBank accession number: GCA 000002985.3), *Caenorhabditis nigoni* (GenBank accession number: GCA 002742825.1), *Caenorhabditis remanei* (GenBank accession number: GCA 001643735.4) and *Caenorhabditis tropicalis* (GenBank accession number: GCA 016735795.1).

The master bait set list was then used to harvest UCEs from 199 genomes available for Rhabditidae through NCBI for the genera *Oscheius*, *Diploscapter*, *Auanema*, and *Caenorhabditis* (supplementary table 12).

#### Phylogenetic reconstruction of *Caenorhabditis*

To test for the accuracy of phylogenetic reconstructions within Rhabditidae using ultraconserved elements compared to previously reported methods (e.g. orthologous genes), a phylogeny for *Caenorhabditis* species and Rhabditidae species were reconstructed. A data matrix of 75% occupancy was created using phyluce align get only loci with min taxa (each UCE that was present in at least 75% of the taxa analyzed). The alignments were concatenated using phyluce align concatenate alignments. The resulting alignment was used as input for IQ-TREE (v 2.3.6), and a maximum likelihood inference was obtained (1000 ultrafast bootstrapping) using the Model Finder Plus feature for selecting the best substitution model for the dataset. The resulting phylogenetic tree was compared to (Stevens et al., 2019) phylogenetic reconstruction through a tanglegram obtained using ape (v 5.8-1) (Paradis & Schliep, 2019) and phytools (v 2.4-4) (Revell, 2024).

### 5.3 Classification model with Ultraconserved elements (UCEs)

#### 5.3.1 UCE data preparation and preprocessing

To analyze the taxonomic signal capture by UCEs, a multi-step workflow was implemented. First, UCE identifiers were extracted from species-specific FASTA files for nematodes within the families Panagrolaimidae and Rhabditidae. These identifiers were then used to construct binary presence-absence matrices, indicating whether each UCE was detected in each strain. It is essential to note that, although we refer to species throughout this section, the dataset comprises strains, with some species represented by multiple strains. Once the presence-absence matrix was constructed, an exploratory data analysis (EDA) was conducted to examine the patterns of the UCE presence across the samples, including the number of UCEs shared among genera and the variability within genera.

To facilitate subsequent machine learning analyses, strains were assigned to their respective genera. This taxonomic assignment was performed using accession numbers retrieved from the European Nucleotide Archive (ENA). The final output consists of genus-level UCE presence-absence matrices, which are the foundation for predictive modeling. For a detailed description of the extraction steps, matrix construction, and taxonomic assignment, refer to Supplementary Information (A.3), where we provide full methodological details, dataset summaries, and the associated scripts.

The data sets used for training and evaluation were derived from genus-level UCE presence-absence matrices, where genera with insufficient representation (frequency ≤ 1) and rows with missing values were removed. After filtering, the Rhabditidae dataset consisted of 197 samples, 8,336 features (UCEs), and four genera: *Oscheius*, *Diploscapter*, *Auanema*, and *Caenorhabditis*. For Panagrolaimidae, 1,595 predictors (UCEs) were analyzed. Following the removal of genera with insufficient representation and rows with missing values, the Panagrolaimidae dataset (including the outgroup from Cephalobidae-*Acrobeloides*) contained 49 samples and five genera: *Acrobeloides*, *Halicephalobus*, *Panagrellus*, *Panagrolaimus*, and *Propanagrolaimus*. In both datasets, UCE columns were used as predictors (features), and the genus column was converted to a factor for classification, see table 1.

**Table 1:**
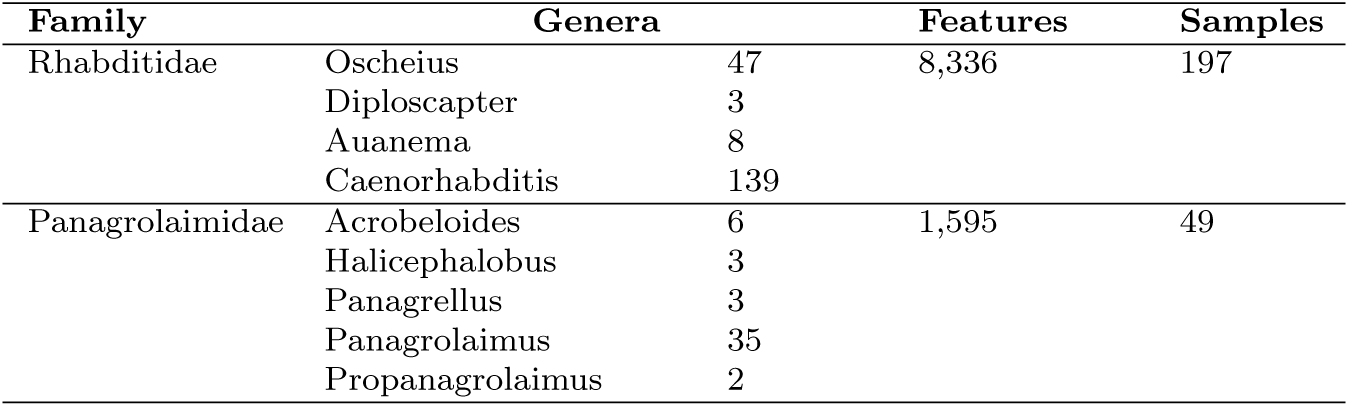
Summary of UCE datasets for machine learning.

#### 5.3.2 Model training, evaluation, and feature selection

The Rhabditidae dataset, which contained the largest number of features and entries, was used to evaluate four machine learning models in R: Random Forest (RF), Logistic Regression (LR), k-Nearest Neighbors (k-NN), and Extreme Gradient Boosting (XGBoost). A stratified partitioning strategy, implemented using the caret package (M. Kuhn, 2008), was employed to divide the data into training (65%) and testing (35%) subsets, preserving the original distribution of genera classes in both sets. Class weights were calculated and incorporated into the Random Forest (RF) and XGBoost models to address class imbalance. Model training utilized five-fold cross-validation, with performance metrics calculated using the multiClassSummary function from the caret package. Random Forest was trained by optimizing the mtry parameter with class weights applied, using the randomForest package (Liaw & Wiener, 2002). Logistic Regression with PCA involved dimensionality reduction through PCA, followed by feature scaling and regularization via the glmnet package (Friedman, Hastie, & Tibshirani, 2010). k-Nearest Neighbors was implemented using default parameters, using the class package (Venables & Ripley, 2002). Extreme Gradient Boosting was trained with class weights incorporated into the learning process, using the xgboost package (Chen & Guestrin, 2016). Performance metrics, including overall accuracy, kappa statistics, sensitivity, and specificity, were used to compare model performance. A confusion matrix provided detailed insight into classification accuracy for each genus. To identify the most relevant features for nematode genera classification, we employed a feature selection approach based on the importance scores generated by the XGBoost model using the xgboost package (Chen & Guestrin, 2016). Specifically, we retained only those features that exhibited a non-zero “Overall” importance score, as determined by the varImp function in the caret package (M. Kuhn, 2008).

#### 5.3.3 Optimized model application to Panagrolaimidae data

To evaluate the performance of the feature-selected XGBoost model in classifying nematode genera within the Panagrolaimidae family, we applied it to the corresponding UCE presence-absence dataset. The “Genera” column was converted to a factor, and genera represented by single occurrences were removed. Missing values were handled using “na.omit()”. The resulting dataset, which included the following sample sizes per genus: *Acrobeloides* (6), *Halicephalobus* (3), *Panagrellus* (3), *Panagrolaimus* (35), and *Propanagrolaimus* (2), exhibited class imbalance (see Table 1). To address this, a stratified split of 65% training and 35% testing was performed using createDataPartition from the caret package (M. Kuhn, 2008), with a fixed seed and manual adjustment to ensure proportional representation. Class weights, inversely proportional to genus frequency, were applied. The XGBoost model was trained using the train function from caret with the xgbTree method, employing 5-fold cross-validation (cv, number = 5) and multiClassSummary for performance evaluation. The Area Under the Curve (AUC) was used as the primary performance metric (metric = “AUC”). Subsequently, the model was retrained using only the most important features, identified by an overall importance value greater than 0, to reduce dataset dimensionality. The test set AUC was then calculated using multiclass.roc from the pROC package (Robin et al., 2011). A confusion matrix was generated to visualize the model’s classification performance.

## 6 Results

### 6.1 Bait design

The bait set design for Panagrolaimidae was conducted by testing eight distinct base genomes and evaluating the number of conserved loci that were shared between one and nine taxa (the eight *Panagrolaimus* taxa from the base genomes and an outgroup of the *Propanagrolaimus* genus). The number of loci targeted across different bait sets varied from 0 to 12,740. To ensure a balance between taxonomic representation and the number of loci targeted, only bait sets that included more than 1,000 loci shared among six or seven taxa were selected for further analysis. In these cases, the number of targeted loci ranged from 1,096 to 2,080, with the corresponding number of probes ranging from 14,018 to 24,217 prior to duplicate removal. After duplicates were removed, the final probe count ranged from 13,565 to 24,006. The Panagrolaimidae bait set was tested on the Rhabditidae family; however, no significant amount of loci could be retrieved. Therefore, a family-specific bait set was designed for Rhabditidae. The preferred bait set for synthesis ultimately targeted 1,612 loci using 18,789 probes.

The bait set designed for Rhabditidae was designed using the *Caenorhabditis elegans* Bristol N2 genome as a base. For the test bait set, six *Caenorhabditis* species were used. The preferred bait set in this case targets 10,397 loci shared among the six datasets used. The number of probes prior to duplicate removal was 124,612, while the final probe count was 121,966.

### 6.2 In-vitro and in-silico testing Panagrolaimidae

SPAdes assemblies of target capture data ranged between 4,2479 and 28,7752 contigs (mean 88,628.05 ±56,482.50529) for Panagrolaimidae exemplars. The two assemblies from the outgroup specimens (Cephalobidae) were more fragmented than in the Panagrolaimidae exemplars (1,121,821 and 1,237,255 contigs). In general, the mean length of the contigs ranged between 215.19 and 1639.88 base pairs (mean 531.86 ± 269.55).

In total, 51 data sets were analyzed and used to reconstruct the phylogeny of the Panagrolaimidae family, including data sets from the outgroup family Cephalobidae (*Acrobeloides*). In total, 1,572 loci with 694,995 informative sites were used, a mean of 442.11 sites per locus, a 95% confidence interval of 8.11, a minimum of 0 sites and a maximum of 1593 sites, with 396 alignments out of 1572 containing more than 0.6 proportion of taxa (n = 30) (alignments can be found in the Zenodo repository – 10.5281/zenodo.15395838).

In the Panagrolaimidae analysis, a total of 49 to 84 ultraconserved elements (UCEs) were retrieved from the outgroup and utilized for phylogenomic analysis, regardless of the data source (target capture or available genome assemblies). Within the genus *Panagrolaimus*, the number of UCEs retrieved and incorporated into the phylogenomic analysis ranged from 479 to 1,457. Specifically, in the target capture dataset, 66 to 80 UCEs were captured in the outgroup, while 277 to 1,200 UCEs were recovered for *Panagrolaimus*. There were 84 UCEs commonly found between all genera analyzed, and 500 UCEs uniquely found in *Panagrolaimus*.

### 6.3 Morphological and molecular evidence for species reassessment

Observations of adult individuals, including both females and males of a recent culture of the strain previously referred to as *Panagrolaimus detritophagus* BSS8, revealed that the nematode originally identified as *Panagrolaimus* “population II” in Boström (1988) and subsequently as *Panagrolaimus detritophagus* BSS8 and *Panagrolaimus* BSS8 is in fact *Neocephalobus halophilus* Paetzold, 1958. The main diagnostic feature of *Neocephalobus*, comparing to *Panagrolaimus*, is the presence of distinct papilliform precloacal sensillum in males and its location some distance anterior from the cloacal opening (Bhat et al., 2025). Additional distinct morphometric features include a longer tail in both males and females in *Neocephalobus* when compared to *Panagrolaimus*, greater pharyngeal region length in *Neocephalobus*, and greater body length body width ratio at mid-body in *Neocephalobus* when compared to *Panagrolaimus* (Supplementary tables **??** and **??** and supplementary text **Morphological description of *Neocephalobus halophilus* BSS8 strain, previously referred to as *Panagrolaimus detritophagus* BSS8**). Furthermore, based on UCEs it is placed as a close relative to *Halicephalobus* species and *Propanagrolaimus* rather than *Panagrolaimus* with a bootstrap support value of 100 (figure 2), in agreement with recently published single-gene phylogenies based on several markers (Bhat et al., 2025).

**Figure 1:**
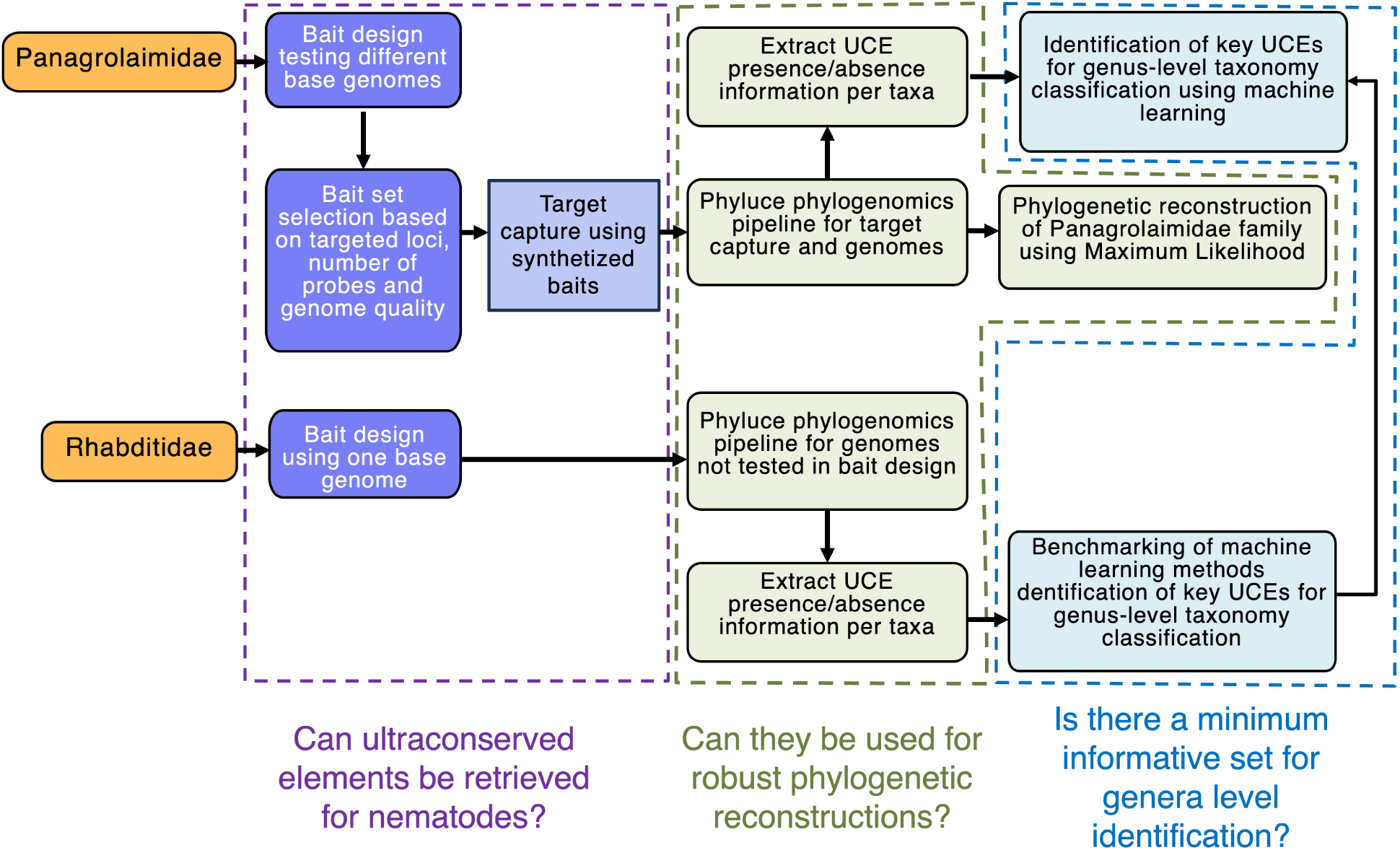
Overview of the workflow implemented in this study. Bait sets were designed and tested for both the Panagrolaimidae and Rhabditidae families. From UCEs harvested in the different families with various approaches, a presence/abscence matrix was obtained to train and test machine learning classifiers to identify the most important UCEs for genus-level classification. For Panagrolaimidae, the model was further tested using UCE data extracted from genome skims generated with Nanopore sequencing. Graph generated based on Canva template from elversa.

**Figure 2:**
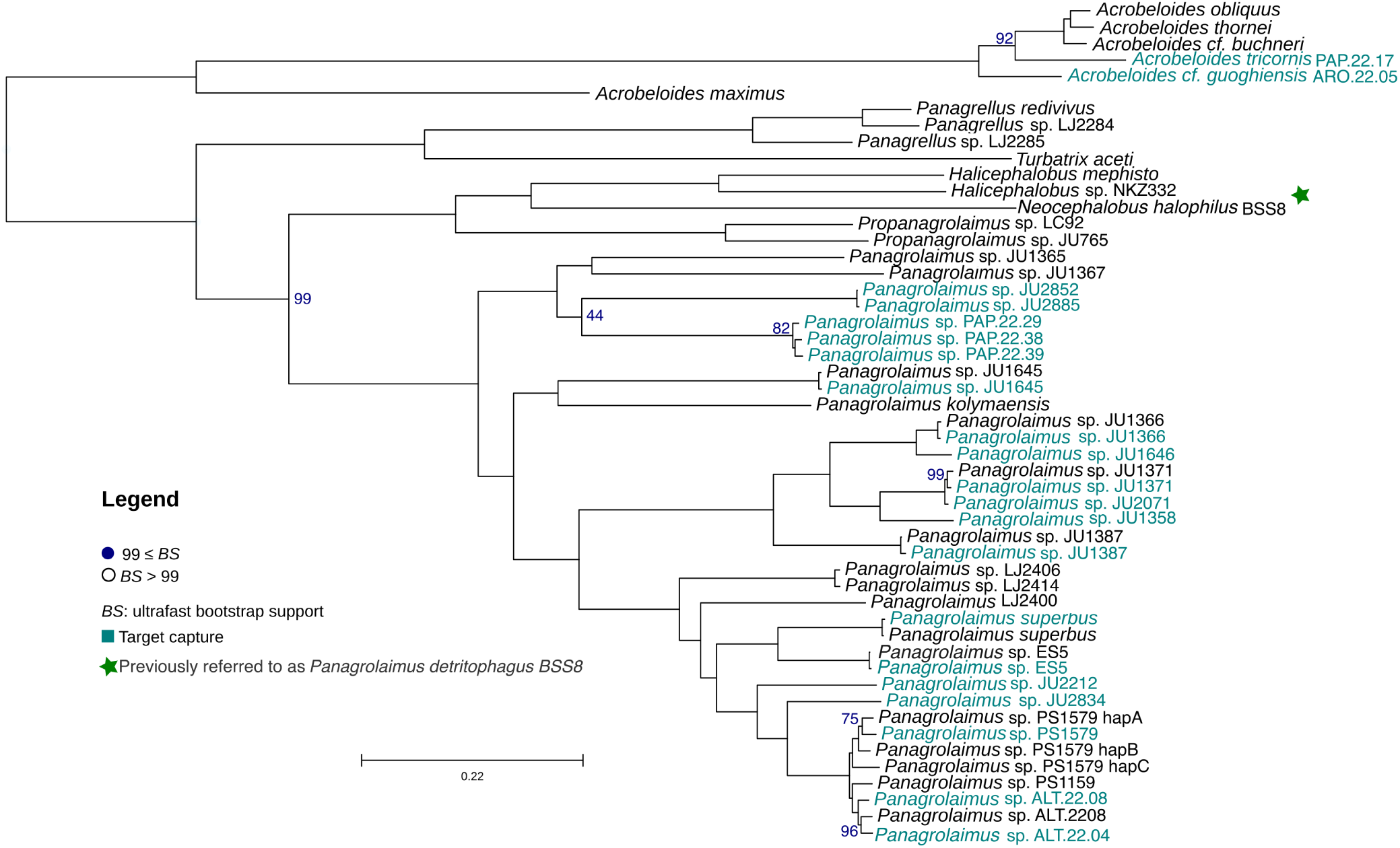
Phylogenetic reconstruction of Panagrolaimidae (*Acrobeloides* is an outgroup) using maximum likelihood, 51 taxa are included in the analysis. Ultraconserved elements (UCEs) from the target capture approach are highlighted in aquamarine, while UCE data harvested from available genomes are highlighted in black. The phylogeny is based on 396 alignments. Only bootstrap values below 99 are shown in gray and orange.

### 6.4 In-silico testing Rhabditidae

The bait set designed with *Caenorhabditis* genomes was then tested on several genera within the Rhabditidae family (*Oscheius*, *Diploscapter*, *Auanema*, and *Caenorhabditis*). The number of UCE loci harvested from the genomes ranged from 1 (*Oscheius myriophilus*) to 5,700 (*Caenorhabditis elegans*). Only 50 UCEs were shared between all genera of the family, with most UCEs being uniquely found in *Caenorhabditis*. A phylogenetic reconstruction with high support was obtained for 64 taxa of the Rhabditidae family (supplementary figure 11).

In total, 27 data sets were analyzed and used to reconstruct the phylogeny of the genus *Caenorhabditis*. Altogether, 945 loci with 437,091 informative sites were used, a mean of 462.53 sites per locus, a 95% confidence interval of 10.03, a minimum of 33 sites and a maximum of 829 sites, with 945 alignments containing more than 0.75 proportion of taxa (alignments can be found in the Zenodo repository – 10.5281/zenodo.15395838).

### 6.5 Classification model with ultraconserved Elements (UCEs)

#### 6.5.1 Presence and characteristics of UCEs in Rhabditidae and Panagrolaimidae

We analyzed the total number of UCEs per strain within the families Rhabditidae and Panagrolaimidae to assess their presence across strains. The results, visualized in the Supplemental Material (Figure 12 and 13), indicate substantial variation in UCE counts between strains within each family. Although most strains exhibit comparable UCE numbers, some show significantly lower counts. Specifically, in Rhabditidae, UCE counts range from 26 to 11,400, with a median of 7,762, while in Panagrolaimidae, they range from 15 to 1,457, with a median of 767.5. This variability may reflect differences in genome characteristics, such as assembly completeness or lineage-specific variation in the retention of UCEs. This variability may also stem from the bait set design, which was based only on a few taxa, for instance, for Rhabditidae, the bait set was obtained only by analyzing the genus *Caenorhabditis*, yet later applied to a broader range of lineages within Rhabditidae, including genera such as *Diploscapter* that were not represented in the initial design.

To ensure data quality, we applied an outlier detection approach based on UCE counts. Instead of relying solely on interquartile range (IQR) methods, which can be sensitive to skewed count distributions of UCEs occurrences per strain, we opted for a more robust approach by removing the bottom 1% of strains with the lowest UCE counts. This method is particularly useful for genomic datasets where sequencing depth and assembly completeness vary, ensuring that only strains with sufficiently high UCE representation are retained for subsequent analyses. After this procedure, three strains were identified as outliers and removed, two from Rhabditidae (GCA 002207785.1 and GCA 964036155.1) and two from Panagrolaimidae (GCA 028622995.1 – *Panagrolaimus kolymaensis* prior to splitting contigs, and GCA 963969345.1 *Turbatrix aceti*).

After filtering, we analyzed how frequently each UCE appears across the remaining strains. The UCE occurrence frequency profile for both families is presented (Fig 4), showing how many UCEs are shared across different proportions of strains. In Panagrolaimidae, the distribution is unimodal, with most UCEs appearing in an intermediate range of strains. In contrast, the Rhabditidae distribution exhibits bimodality, indicating two groups of UCEs with different levels of conservation. The low-frequency group (0-50) may represent UCEs that are present in some species but absent in others, reflecting potential lineage-specific losses or variation in presence or absence. Conversely, the high-frequency group (100-150) comprises UCEs that are widely shared across Rhabditidae, suggesting they may be under stronger evolutionary constraints, potentially due to their regulatory or structural functions.

**Figure 3:**
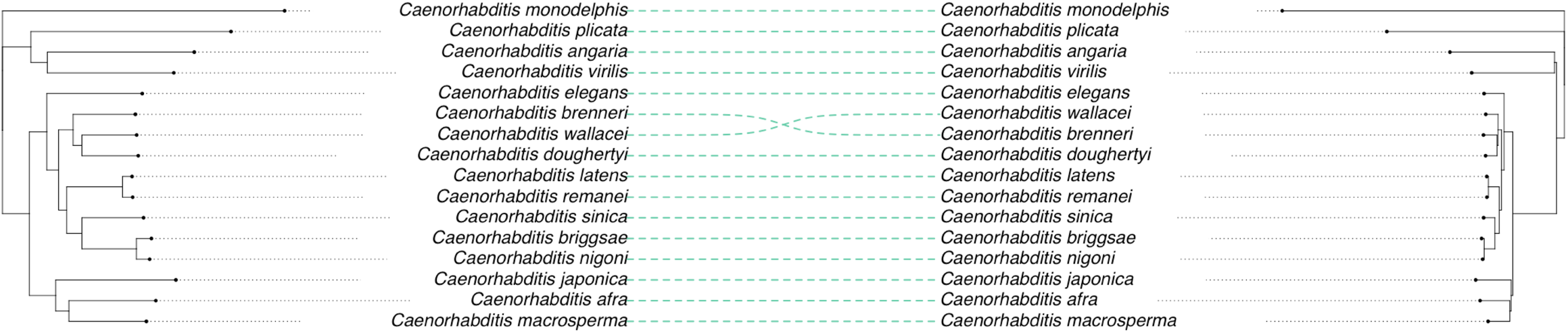
Tanglegram for the genus *Caenorhabditis* comparing UCE-derived reconstruction (left) obtained in this work and gene orthology-based reconstruction (right) from (Stevens et al., 2019). The alternative topology for *C. wallacei* and *C: brenneri* is also proposed in Stevens et al., 2019 depending on the method used.

**Figure 4:**
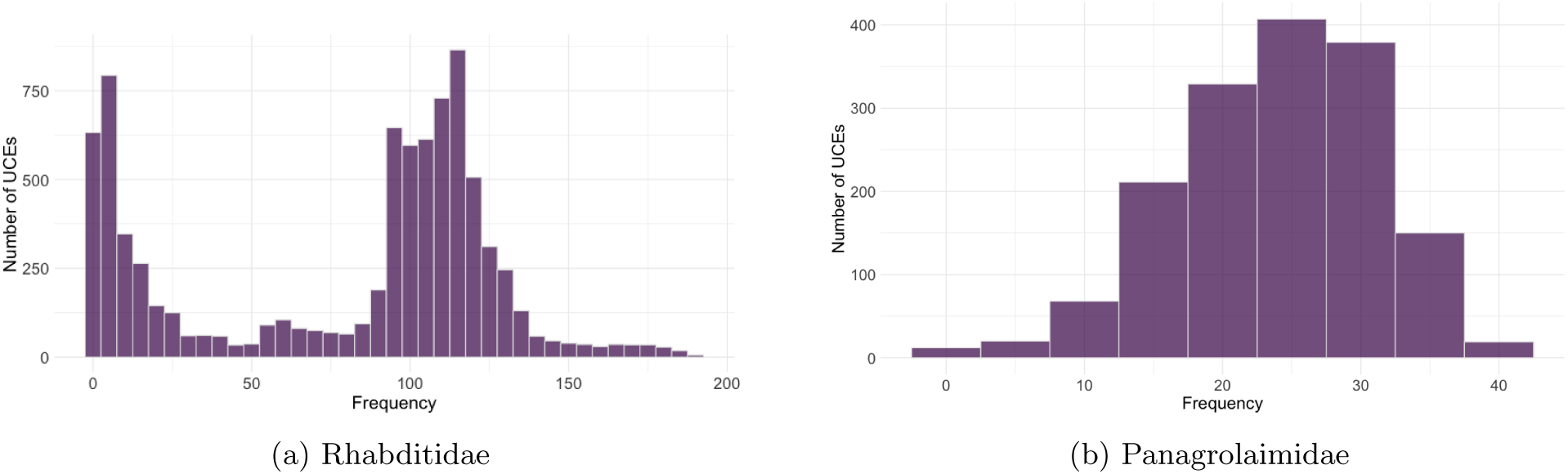
Divergent patterns of UCE occurrence frequency across strains in Rhabditidae and Panagrolaimidae. The x-axis represents the percentage of strains in which a given UCE is present, while the y-axis indicates the number of UCEs observed at each frequency level. The unimodal pattern in Panagrolaimidae contrasts with the bimodal pattern in Rhabditidae, suggesting differences in evolutionary constraints and genomic conservation across the two families.

We analyzed the number of UCEs shared across genera within each family. In Rhabditidae, a total of 50 UCEs are shared across the genera *Auanema*, *Caenorhabditis*, *Diploscapter*, and *Oscheius* (Fig 5a). In Panagrolaimidae, 84 UCEs are shared across the genera *Panagrolaimus*, *Halicephalobus*, *Panagrellus*, *Propanagrolaimus* and the outgroup genus *Acrobeloides* (Fig 5b). The corresponding Venn diagrams summarizing these overlaps are presented in Fig 5, and the full list of shared UCEs is available in Supplementary Table 13

**Figure 5:**
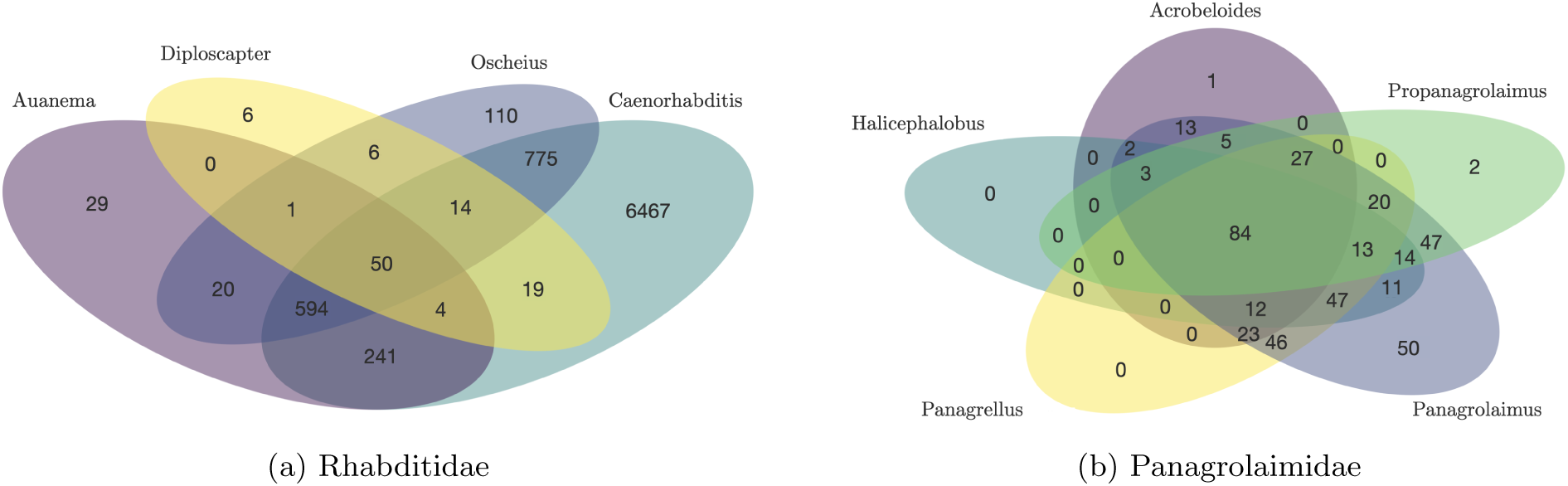
Venn diagrams for shared UCEs across genera in Rhabditidae and Panagrolaimidae. The diagrams illustrate the number of UCEs shared across the genera within each family.

#### 6.5.2 Benchmark of machine learning models on Rhabditidae data

The performance evaluation of the four machine learning models: Random Forest (RF), Logistic Regression (LR) with data transformed with PCA, k-Nearest Neighbors (k-NN), and Extreme Gradient Boosting (XG-Boost) on the Rhabditidae dataset of UCEs, along with their respective performance metrics, is shown in Table 2. A comparative analysis revealed differences in classification performance among the models. XGBoost achieved the highest AUC score of 0.9997, demonstrating superior predictive power. It correctly classified all *Auanema* and *Diploscapter* samples and misclassified only one *Oscheius* sample as *Caenorhabditis*. Its strong performance is attributed to its gradient boosting framework and the integration of class weights, which effectively handled class imbalance. Random Forest performed well, achieving an AUC of 0.9939, but struggled with *Oscheius* and *Diploscapter*. Specifically, it misclassified nine out of sixteen *Oscheius* samples as *Diploscapter*, leading to reduced accuracy in predicting minority classes. While class weighting helped mitigate imbalance, it was not as effective as XGBoost’s approach. k-Nearest Neighbors (k-NN) obtained an AUC of 0.9858 and showed difficulty in classifying *Oscheius* and *Diploscapter*. Two *Caenorhabditis* samples were misclassified as *Oscheius*, and one *Oscheius* sample was wrongly predicted as *Diploscapter*, indicating sensitivity to the choice of k and the distance metric in high-dimensional data. Logistic Regression with PCA had the lowest performance, with an AUC of 0.8290. While PCA successfully reduced dimensionality and improved computational efficiency, the model struggled with minority class classification. It failed to predict *Auanema* and *Diploscapter* entirely and misclassified multiple *Oscheius* samples, demonstrating limitations despite regularization techniques to prevent overfitting. Overall, XGBoost emerged as the most effective model, demonstrating the highest classification accuracy and robustness across all classes. Its ability to handle imbalanced datasets, coupled with its efficiency and strong generalization capacity, makes it particularly suited for classifying genera within Rhabditidae. The complete confusion matrices and class-specific performance metrics for each model are provided in the Supplementary Materials A.4.

**Table 2:** Performance metrics for machine learning models on the Rhabditidae family.

| Model | AUC | Accuracy (%) | Kappa | Sensitivity (%) | Specificity (%) | Best hyperparameter |
| --- | --- | --- | --- | --- | --- | --- |
| RF | 0.9939 | 85.07 | 0.6819 | 85.42 | 96.21 | mtry = 3 |
| LR with PCA | 0.8290 | 94.03 | 0.8599 | 74.46 | 98.55 | PCA-transformed |
| k-NN | 0.9858 | 94.03 | 0.8652 | 73.96 | 97.68 | k = 5 |
| XGBoost | 0.9997 | 98.51 | 0.9657 | 99.48 | 99.52 | Depth = 6, Learning rate = 0.3 |

##### Identification of important features UCEs

Following the application of the non-zero importance score feature selection method on the XGBoost model, a total of 46 features were identified as relevant for nematode genera classification on the Rhabditidae family. These features, along with their corresponding “Overall” importance scores, are detailed in the Supplementary Material Table 14. To provide a concise overview of the feature importance, the top 20 features ranked by their “Overall” importance scores are presented in Supplementary Figure 16. The feature uce.15118 exhibited the highest importance score (100.00), indicating its central contribution to the model’s predictive accuracy. Notably, the top six features (uce.15118, uce.1361, uce.2297, uce.6206, uce.17604, uce.16976) demonstrated significantly higher importance scores compared to the remaining features, suggesting their critical role in discriminating between nematode genera for the family Rhabditidae. Furthermore, six features were also found to be crucial in the cross-validation with varying features, where the model’s AUC significantly improved when these features were included.

#### 3.5.3 Classification performance on Panagrolaimidae UCE data

The optimized XGBoost model worked well on the Panagrolaimidae UCE dataset. To address the class imbalance, stratified sampling and weighted training were applied. The model was first trained on all UCE features, and next, the training was done using just the top 39 most important UCEs (i.e., those with an overall importance value *>*1, see Supplemental Material Table 15). Both the full model and the reduced feature model achieved the same overall accuracy of 94.12%. However, there were slight differences in the confusion matrices and other classification metrics (Supplementary material A.5). These results suggest that the reduced set of features retained the model’s predictive power while reducing dimensionality. Detailed statistics and metrics per class are provided in the supplementary material (Section A.5).

## 7 Discussion

### 7.1 Ultraconserved elements can be identified and retrieved for nematodes and can be used for robust phylogenetic reconstructions

We designed two bait sets to capture ultraconserved elements within the phylum Nematoda, with a focus on the families Rhabditidae and Panagrolaimidae. We then identified the most informative UCEs using machine learning classifiers to further aid in nematode classification, utilizing genetic information at high resolution. We demonstrate that using the target-captured loci, a well-resolved phylogeny can be obtained for Panagrolamidae, placing correctly described taxa within the family and showing high support values for most nodes, providing insights into the phylogenetic relationships among taxa that previously had no genetic data available.

Our phylogenetic analysis is consistent with previous studies based on both molecular markers and wholegenome data (Abolafia & Vecchi, 2021; Qing et al., 2024; Shokoohi & Masoko, 2024), with the exception of *Panagrolaimus* BSS8, which we now confirm with both molecular and morphological evidence, that it is in fact *Necephalobus halophilus*, being closely related to the genus *Halicephalobus* in our phylogenetic reconstruction. The nematode genus *Panagrolaimus* appears to be more derived within the Panagrolaimidae family, and it is resolved as the closest relative to the genus *Propanagrolaimus*. Both genera form a monophyletic clade that is closely related to *Halicephalobus*, which in turn is sister to *Panagrellus*. All of these genera are distinct from the outgroup, *Acrobeloides*, which is placed outside of the Panagrolaimidae clade.

We also find asexual Panagrolaimus clustering together (*Panagrolaimus sp.* PS1579, *Panagrolaimus sp.* PS1159 Lewis et al. (2009) and the newly isolated *Panagrolaimus sp.* ALT.22.08) (Villegas et al., 2024)) and separately from sexually reproducing ones (*Panagrolaimus einhardii* and *Panagrolaimus superbus* Lewis et al. (2009)). Furthermore, the asexual nematode *Panagrolaimus kolymanesis*, previously placed as an outgroup to all other *Panagrolaimus* nematodes Shatilovich et al. (2023), is here shown to cluster with *Panagrolaimus JU1645* and does not sit on a branch of its own. We find with high support that sexually reproducing strains from the Atacama Desert Villegas et al. (2024) cluster together and are more closely related to other basal *Panagrolaimus* strains as *Panagrolaimus sp.* JU1367 and *Panagrolaimus sp.* JU1365.

In the Rhabditidae family, we see that our phylogenetic reconstruction using UCEs is congruent when compared to previous work (Stevens et al., 2019), showing a clear distinction between the *Elegans* and *Japonica* groups and both forming the *Elegans* supergroup with *Caenorhabditis monodelphis* as an outgroup. While *C. brenneri* and *C. wallacei* do not exhibit a direct one-to-one correspondence between their placement in the phylogenetic reconstruction, this alternative hypothesis remains consistent with the phylogenetic trees reported in (Stevens et al., 2019), which were generated using different methods.

While using UCEs in both families is a promising approach for phylogenetic reconstructions and further analysis, it is important to note that these “ultraconserved” elements often exhibit conservation only within relatively narrow taxonomic scales (e.g., within families rather than across orders). This pattern suggests a higher rate of sequence divergence across broader nematode lineages, consistent with the rapid molecular evolution observed in the phylum.

### 7.2 *Panagrolaimus detritophagus* BSS8 is in fact *Neocephalobus halophilus* BSS8 Paetzold, 1958

The *Panagrolaimus* BSS8 strain is one of the cultures of nematodes established by Dr. Björn Sohlenius from samples from the Surtsey Island (Lewis et al., 2009). There were three different populations of Panagrolaimus collected on Surtsey, established in cultures and described by (Bostrom, 1988), of which the *Panagrolaimus* “population II” matches the morphology of BSS8 strain, in both morphology and in measurements, but especially in the presence of distinct midventral precloacal papilliform sensillum, located at a distance from cloacal opening, at level with spicule manubrium. The population was never identified to species level by (Bostrom, 1988). Still, it was considered to be most similar to *Panagrolaimus chalcographi* by using the key from (Andŕassy, 1984) or *Panagrolaimus detritophagus* by using the (Williams, 1986) grouping. The conclusion of the manuscript was that the population II from Surtsey “appears intermediate between *Panagrolaimus detritophagus* and *Panagrolaimus chalcographi*” (Boström, 1988) but no conclusive decision was made. In subsequent publications, this culture was named (without any obvious justification) as *Panagrolaimus detritophagus* BSS8 (Felix et al., 2000; Lewis et al., 2009; Schiffer et al., 2014; Shannon, Browne, Boyd, Fitzpatrick, & Burnell, 2005). Here we provide additional morphological evidence to confirm the taxonomic identity of the BSS8 strain.

Presence of distinct midventral precloacal papilliform sensillum located at a distance from cloacal opening (at the level of spicule manubrium) is one distinct feature of this population, separating it from most known species currently classified in the genus *Panagrolaimus* and all currently accepted genera of Panagrolaimidae. Only the following three species in Panagrolaimidae are morphologically similar to the BSS8 strain:

*Neocephalobus aberrans* Steiner, 1929, originally described as Cephalobus (Neocephalobus) aberrans by (Steiner, 1929b)) and later transferred to the genus *Panagrolaimus* (= *Panagrolaimus aberrans* (Steiner, 1929b) Godey, 1963) was found in the feces of a guinea pig, kept at the University of California. No measurements were included in the original description, which is rather incomplete in general. However, type population of *Neocephalobus aberrans* can be easily distinguished from the BSS8 in having narrow long stoma with minute tooth, whereas stoma in BSS8 is broad and the tooth is distinct. Moreover, excretory pore in *N. aberrans* is surrounded by a cuticularised ring (absent in BSS8) and shorter tail in both females and in males. The newly discovered population of the same species (Bhat et al., 2025) can also be clearly separated from the BSS8 strain in having longer and narrower stoma (just like in the original description), presence of median bulb, shorter tail and different arrangement of male caudal sensilla.

*Neocephalobus halophilus* Paetzold, 1958, also later transferred to the genus *Panagrolaimus* and renamed to *Panagrolaimus paetzoldi* Goodey, 1963 (homonym of *Panagrolaimus halophilus* Meyl, 1954), was collected in salt marshes in central Germany. Morphologically, it matches the BSS8 strain in all parameters, including the shape of stoma, number and arrangement of sensory structures on male tail Bhat et al. did not make any taxonomic decisions on the status of the BSS8 strain, however, they positively identified Dutch populations of Panagrolaimus paetzoldi as Neocephalobus halophilus, based solely on the molecular data. These populations form a well supported clade with BSS8 but are different by about 24 substitutions. Unfortunately, there is no morphological voucher data available for these sequences. Here we propose to consider them a distinct population of Neocephalobus halophilus until new specimens are collected and sequenced.

*Panagrolaimus orthomici* Korentchenko, 1992 (Korentchenko, 1992) can be easily separated from BSS8 in having two subdorsal teeth in the stoma (single in BSS8), more anterior position of the excretory pore, different arrangement of sensory structures on male tail. This species is here transferred to the genus *Neocephalobus*, as *Neocephalobus orthomici* (Korentchenko, 1992) comb. n.

In conclusion, currently available morphological evidence, along with phylogenetic placement based on UCEs, as well as recently published reinstatement of the genus *Neocephalobus* (Bhat et al., 2025), confirms that the nematode originally identified as *Panagrolaimus* “population II” in (Bostrom, 1988) and subsequently as *Panagrolaimus detritophagus* BSS8 and *Panagrolaimus* BSS8 is in fact *Neocephalobus halophilus* Paetzold, 1958.

### 7.3 A minimum UCE informative set can accurately classify taxa to genus level

Applying machine learning to UCE presence-absence data provided valuable insights into taxonomic classification within Nematoda. In Rhabditidae, the XGBoost model outperformed all other models (Random Forest, PCA-Logistic Regression, and k-Nearest Neighbors) with an AUC score of 0.9997. Feature selection using the non-zero importance scores derived from XGBoost revealed that 46 UCEs are important for genus-level classification, with uce.15118 being the most informative. The optimized XGBoost model strategy was applied to the Panagrolaimidae dataset, showing its versatility. Both the full model (with the complete set of features) and the model using only the top 39 most important UCEs (importance *>* 1) reached the same overall accuracy of 94.12%. This shows that an efficient classifier for taxonomic assignment can be constructed from a smaller set of informative UCEs.

Through comparative analyses, it was observed that certain UCEs are shared across genera. The most frequently observed UCEs were also important in the XGBoost model. In Panagrolaimidae, the uce.9910, uce.194, uce.8674, uce.459, uce.8786, uce.323 appeared in all three categories, indicating their potential role in genus differentiation. Although XGBoost can predict well, it is essential to recognize its limitations. The performance of the model people create will depend upon the quality and completeness of the input data, and any species sampling biases or gaps in the UCE dataset can affect results.

### 7.4 UCE-Based Nematode Classification potential for biodiversity assessments and agricultural monitoring

Nematodes serve as essential bioindicators of soil quality and ecosystem function, making their rapid and accurate classification essential for environmental monitoring and agricultural research. Their classification in colonizer-persister categories (cp) as well as the characterization of feeding types based on taxonomic classification (family and genera respectively) have been used for soil health characterization (Du Preez et al., 2022; Ferris, Bongers, & de Goede, 2001; T. Martin, Wade, Singh, & Sprunger, 2022). For instance, opportunistic nematodes (cp-1 and cp-2) reproduce fast and indicate disturbed, nutrient-enriched, or polluted soils, whereas persisters (cp-4 and cp-5) reproduce slowly and indicate stable ecosystems with good soil health. Additionally, the types of feeding and the proportion of different feeding types in soils can be indicative of high organic input, organic matter decomposition, or potential nutrient imbalances. Their rapid identification can enable accurate, near-real-time biodiversity assessments, for instance, through the use of portable sequencing technologies (e.g., Nanopore) and the harvesting of UCEs, as well as model classification. Recent studies have demonstrated that whole genome amplification techniques (e.g., MDA) can generate sufficient DNA from single nematode individuals for successful Nanopore sequencing (Lee et al., 2023; Roberts, Gilmore, Struck, & Kocot, 2024). This opens up the possibility of combining UCE-based approaches with single-worm sequencing, allowing for accurate taxonomic identification and downstream analyses directly on-site, even in cases where culturing is challenging or sample availability is limited.

Such methods can be extended beyond free-living nematodes and applied to the analysis of parasitic nematodes, which are currently a critical target for molecular diagnostics due to their significant impact on global agricultural systems and animal health (Abad et al., 2008; Gang & Hallem, 2016; Strydom, Lavan, Torres, & Heaney, 2023). Particularly focusing on agricultural systems, genera as *Meloidogyne*, *Heterodera*, and *Pratylenchus* are known for causing significant crop losses globally (Khan, 2023). Their early detection is vital for timely management strategies (J. T. Jones et al., 2013; Z^̌^ibrat et al., 2023). Integrating molecular tools such as ultra-conserved elements (UCEs) with portable sequencing technologies (e.g., Nanopore) has the potential to substantially accelerate the detection of plant-parasitic nematodes (PPNs) by soil testing facilities and quarantine labs. Notably, the aforementioned PPN genera are closely related to each other (Thapa, Gates, Reuter-Carlson, Androwski, & Schroeder, 2019), and belong to Clade IV of the Nematoda, alongside the families Panagrolaimidae and Cephalobidae, which were analyzed in this study (M. Blaxter, 2011). This phylogenetic proximity suggests that the bait set developed here could also be effective for targeting UCEs from PPN taxa. Consequently, this approach could serve as a valuable tool to advance both ecological research and applied agricultural diagnostics.

Future advancements in automation could further enhance the speed and scalability of taxonomic classification pipelines, making them even more suitable for large-scale ecological and environmental monitoring. Currently, our approach focuses on taxa within Rhabditidae and Panagrolaimidae; however, we anticipate that extending this method to other ecologically relevant nematode taxa could provide valuable insights, particularly in soil health assessments.

Our study highlights the effectiveness of the newly designed bait sets for multiple loci (UCEs) and target enrichment as a valuable genomic resource for investigating evolutionary relationships among nematodes, specifically within the Rhabditidae and Panagrolaimidae families. This approach enables the retrieval of thousands of genetic markers with known homology, producing results comparable to those obtained using orthologous genes, while allowing for the study of a broader range of taxa at lower costs. While not explored here, the high variability of the flanking region in the targeted conserved loci can allow for fine-scale analysis at the population level, enabling comparisons between the same species occurring at different geographical locations or under different environmental stressors.

## 8 Author contributions

The study was designed by L.V., L.J. and P.H.S.. J.v.S provided support with bioinformatic pipelines. Laboratory work was conducted by L.V Bioinformatic and statistical work was conducted by L.V. and L.J. P.H.S., A-M.W and O.H. provided supervision during the entire project. The manuscript draft was written by L.V. and L.J., all authors further edited the manuscript.

## A Supplementary Information

### Annex

#### A.1 Bait set design

**Table 3:**
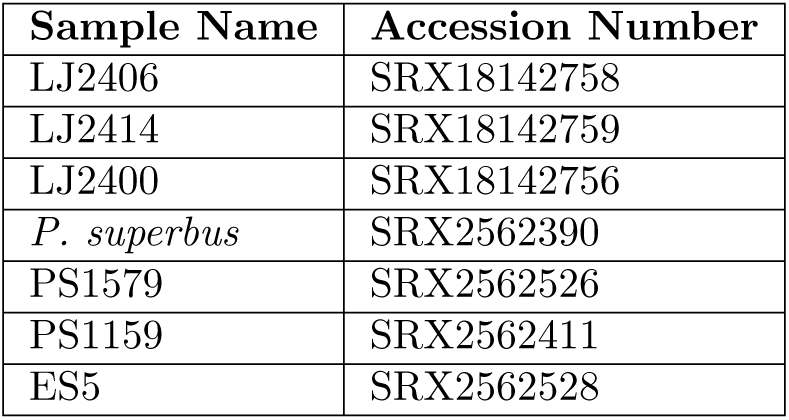
Re-sequencing data accession numbers of the different strains tested for the bait set design of the Panagrolaimidae family.

**Table 4:** Genera and corresponding GenBank accession numbers for phylogenetic reconstruction of Panagrolaimidae family.

| Genera | GenBank Accession |
| --- | --- |
| *Panagrellus* | GCA_341325.1 |
| *Halicephalobus* | GCA_9193035.1 |
| *Halicephalobus* | GCA_9761265.1 |
| *Panagrolaimus* | GCA_28622995.1 |
| *Acrobeloides* | GCA_34698545.1 |
| *Acrobeloides* | GCA_34699885.1 |
| *Acrobeloides* | GCA_34700925.1 |
| *Panagrolaimus* | GCA_963922195.1 |
| *Turbatrix* | GCA_963969345.1 |
| *Panagrolaimus* | GCA_964035985.1 |
| *Propanagrolaimus* | GCA_964059925.1 |
| *Neocephalobus* | GCA_964187885.1 |
| *Acrobeloides* | GCA_964212105.1 |
| *Propanagrolaimus* | GCA_964245515.1 |
| *Panagrolaimus* | GCA_964249645.1 |

**Table 5:** Species and strain names and isolation area of cultures used for generating target capture data in this study.

| Species name or strain code | Isolation Origin |
| --- | --- |
| *Acrobelloides* cf. *guoghiensis* ARO.22.05 | Atacama Desert, Chile |
| *Panagrolaimus* sp. ALT.22.04 | Atacama Desert, Chile |
| *Panagrolaimus* sp. ALT.22.08 | Atacama Desert, Chile |
| *Panagrolaimus* sp. JU2834 | Kelaat Sraghna, Maroc |
| *Panagrolaimus* sp. PS1579 | California, United States |
| *Panagrolaimus superbus* | Surtsey Island, Iceland |
| *Panagrolaimus* sp. JU1358 | Kerala, India |
| *Panagrolaimus* sp. JU1366 | Tamil Nadu, India |
| *Panagrolaimus* sp. JU1371 | Pondicherry, India |
| *Panagrolaimus* sp. JU1387 | La Reunión, France |
| *Panagrolaimus* sp. JU1645 | Santo Antao Island, Cape Verde |
| *Panagrolaimus* sp. JU1646 | Santiago Island, Cape Verde |
| *Panagrolaimus* sp. JU2071 | Europa Island, France |
| *Panagrolaimus* sp. JU2212 | Hüschul uul, Mongolia |
| *Panagrolaimus* sp. JU2852 | Cordoba, Argentina |
| *Panagrolaimus* sp. JU2885 | Lofoten Islands, Norway |
| *Acrobelloides tricornis* PAP.22.17 | Atacama Desert, Chile |
| *Panagrolaimus* sp. PAP.22.29 | Atacama Desert, Chile |
| *Panagrolaimus* sp. PAP.22.38 | Atacama Desert, Chile |
| *Panagrolaimus* sp. PAP.22.39 | Atacama Desert, Chile |

**Table 6:** Species and strains DNA concentration with corresponding 260/280 and 260/230 ratios measured using Nanodrop.

| Strain or species | Concentration (ng/µl) | 260/280 | 260/230 |
| --- | --- | --- | --- |
| *Acrobelloides tricornis* PAP.22.17 | 15 | 1.87 | 0.76 |
| *Acrobelloides* cf. *guoghiensis* ARO.22.05 | 9 | 1.72 | 0.54 |
| *Panagrolaimus* sp. JU2852 | 33.1 | 1.98 | 1.04 |
| *Panagrolaimus* sp. JU2885 | 51.3 | 1.81 | 1.21 |
| *Panagrolaimus* sp. PAP.22.29 | 39.8 | 1.86 | 0.08 |
| *Panagrolaimus* sp. PAP.22.38 | 11.3 | 1.83 | 0.07 |
| *Panagrolaimus* sp. PAP.22.39 | 32.8 | 1.93 | 1.35 |
| *Panagrolaimus* sp. JU1645 | 35.5 | 1.92 | 0.18 |
| *Panagrolaimus* sp. JU1366 | 12.6 | 1.95 | 0.09 |
| *Panagrolaimus* sp. JU1646 | 30.4 | 1.71 | 0.38 |
| *Panagrolaimus* sp. JU1371 | 33.3 | 1.93 | 0.64 |
| *Panagrolaimus* sp. JU2071 | 57.6 | 1.92 | 1.37 |
| *Panagrolaimus* sp. JU1358 | 28.7 | 1.75 | 0.41 |
| *Panagrolaimus* sp. JU1387 | 41.9 | 1.93 | 1.03 |
| *Panagrolaimus superbus* | 78.1 | 1.95 | 1.06 |
| *Panagrolaimus* sp. ES5 | 66.3 | 1.95 | 1.24 |
| *Panagrolaimus* sp. JU2212 | 39.3 | 1.97 | 0.72 |
| *Panagrolaimus* sp. JU2834 | 52.9 | 1.76 | 0.78 |
| *Panagrolaimus* sp. PS1579 | 69.6 | 1.91 | 0.88 |
| *Panagrolaimus* sp. ALT.22.08 | 11 | 1.65 | 0.15 |
| *Panagrolaimus* sp. ALT.22.04 | 66.7 | 1.91 | 2.53 |

#### A.2 Morphological description of *Neocephalobus halophilus* BSS8 strain, previously referred to as *Panagrolaimus detritophagus* BSS8

##### Adult

Body nearly straight when killed by heat, posterior end more curved in males, but not J-shaped. Cuticle finely annulated. Lateral alae with three incisures extending to phasmid. Lip region continuous with body contour; lips partially merged in three pairs, one dorsal and two ventrolateral, although six individual tips are discernible at highest magnification. Labial and cephalic sensilla and amphids are indistinct. Stoma anisomorphic, with distinct metastegostomatal dorsal tooth. Pharynx distinctly subdivided into corpus, isthmus and basal bulb: pharyngeal corpus broad cylindrical; isthmus much narrower, demarcated by a break in muscular tissue; basal bulb oval, with distinct grinder. Nerve ring and deirid at level of isthmus. Excretory pore location varies between base of corpus to anterior part of basal bulb level (supplementary figures 7 and 8).

##### Female

Reproductive system monodelphic, prodelphic, located on the right side of the intestine (dextral); ovary straight. Oviduct very short, about one- half of the corresponding body diameter. Spermatheca axial, well developed. Postvulval uterine sac equal to the corresponding body diameter. Vagina straight, vulval lips protruding. Rectum short. Tail straight, elongate-conoid. Phasmids located at anterior third of tail length (supplementary figures 6 and 9).

##### Male

Reproductive system monorchic, with testis reflexed ventrad. Spicules slender, paired and symmetrical, weakly curved ventrad, with oval manubrium and gradually narrowing shaft. Gubernaculum plate-like. Genital papillae distributed as follows: one pair of subventral precloacal papilliform sensilla located anterior to spicules, single unpaired and much larger precloacal papilliform sensillum located at level of spicule manubrium, one pair of subventral precloacal papilliform sensilla located just anterior to clocal opening, one pair of subventral postcloacal papilliform sensilla located between cloacal opening and phasmid, one pair of subventral postcloacal papilliform sensilla and one pair of subdorsal postcloacal papilliform sensilla located near phasmid, at about the middle of tail length. Tail straight, elongate-conoid (supplementary figure 10).

**Figure 6:**
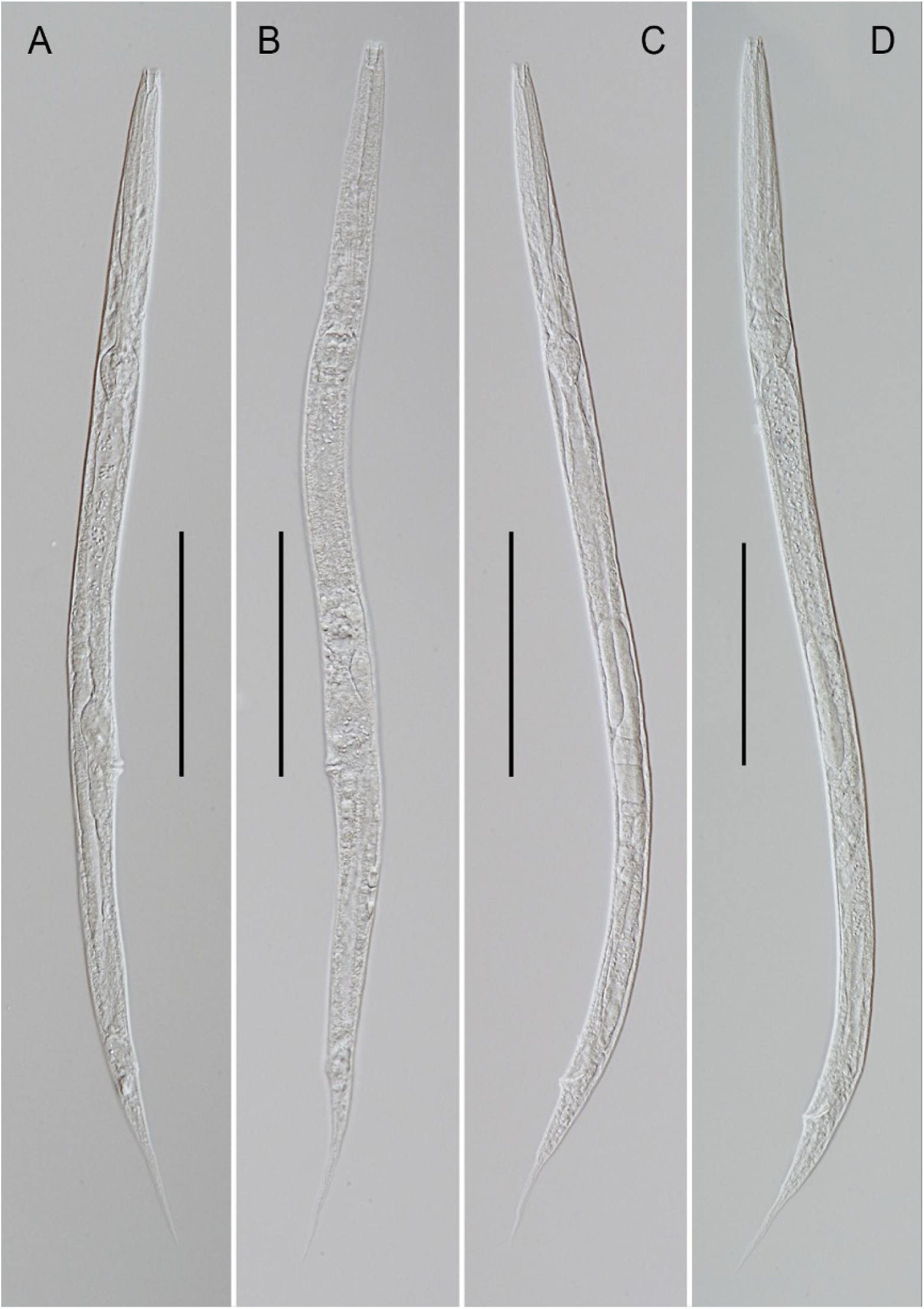
*Neocephalobus halophilus* Paetzold, 1958 (strain BSS8). Entire female (A-B) and male (C-D). Scale bars: A-D = 100 *µ*m.

**Figure 7:**
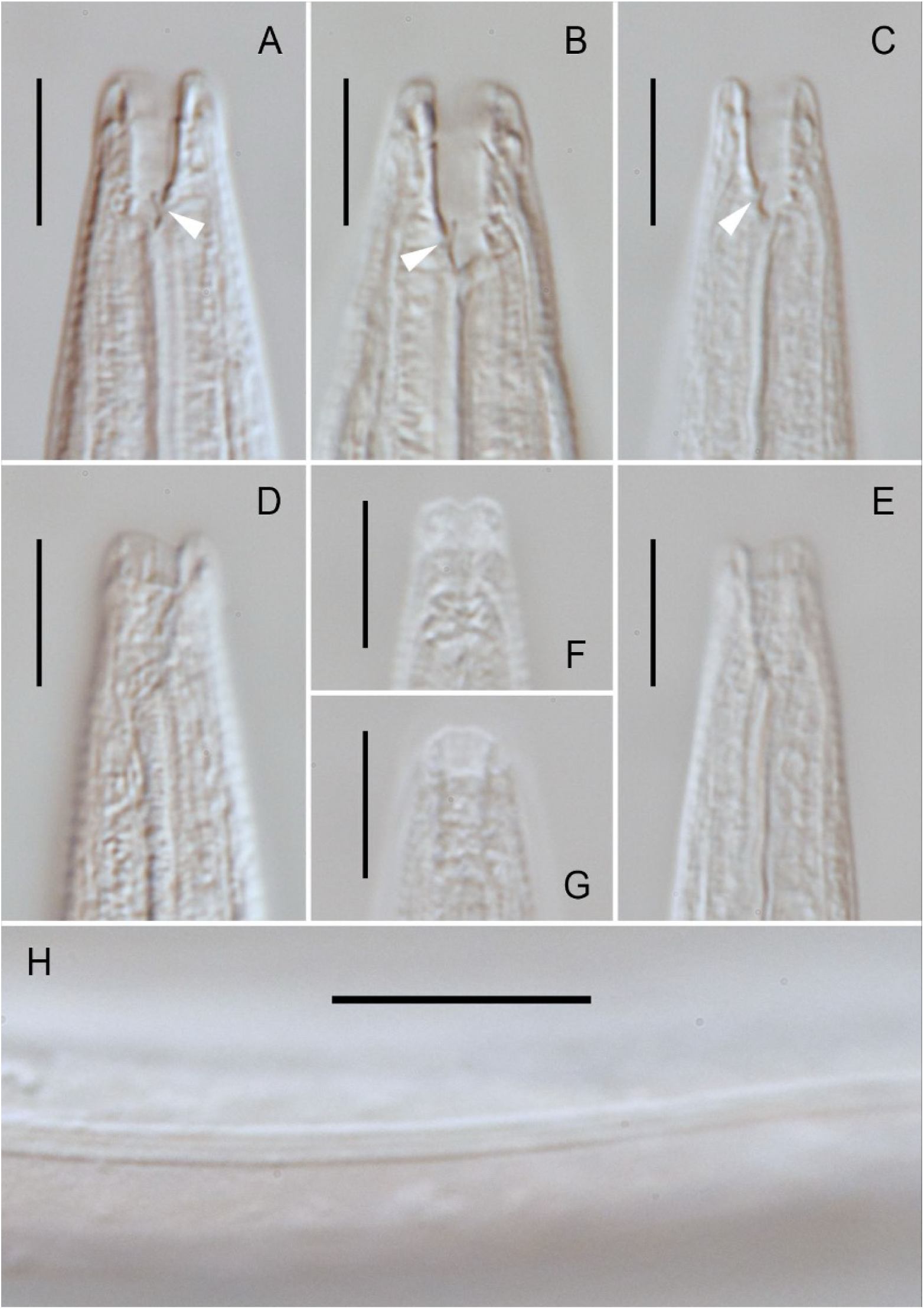
*Neocephalobus halophilus* Paetzold, 1958 (strain BSS8). A-C: Anterior body end, median section (dorsal to the right in A and to the left in B-C, arrow points to the dorsal tooth in A-C); D-E: Lateral view of the labial region (dorsal to the right in D and to the left in E); F: Ventral view of the labial region; G: Dorsal view of the labial region; H: Lateral alae. Scale bars: A-G = 10 *µ*m, H = 20 *µ*m.

**Figure 8:**
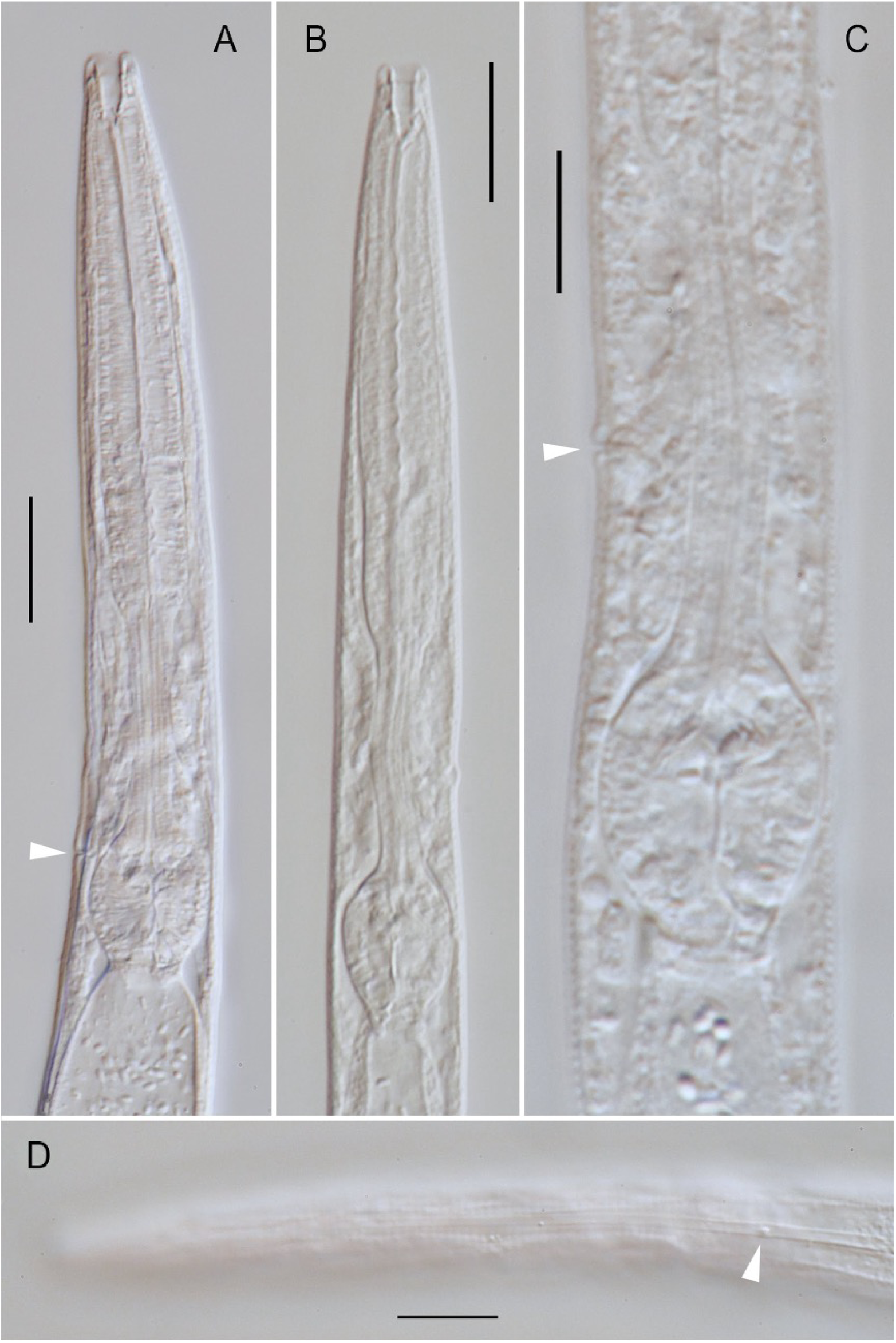
*Neocephalobus halophilus* Paetzold, 1958 (strain BSS8). A-B: Pharyngeal region, median section, showing excretory pore in A (arrow). C: Isthmus and basal bulb, showing excretory pore (arrow). D: Surface view of the pharyngeal region, showing deirid (arrow). Scale bars: A-D = 20 *µ*m

**Figure 9:**
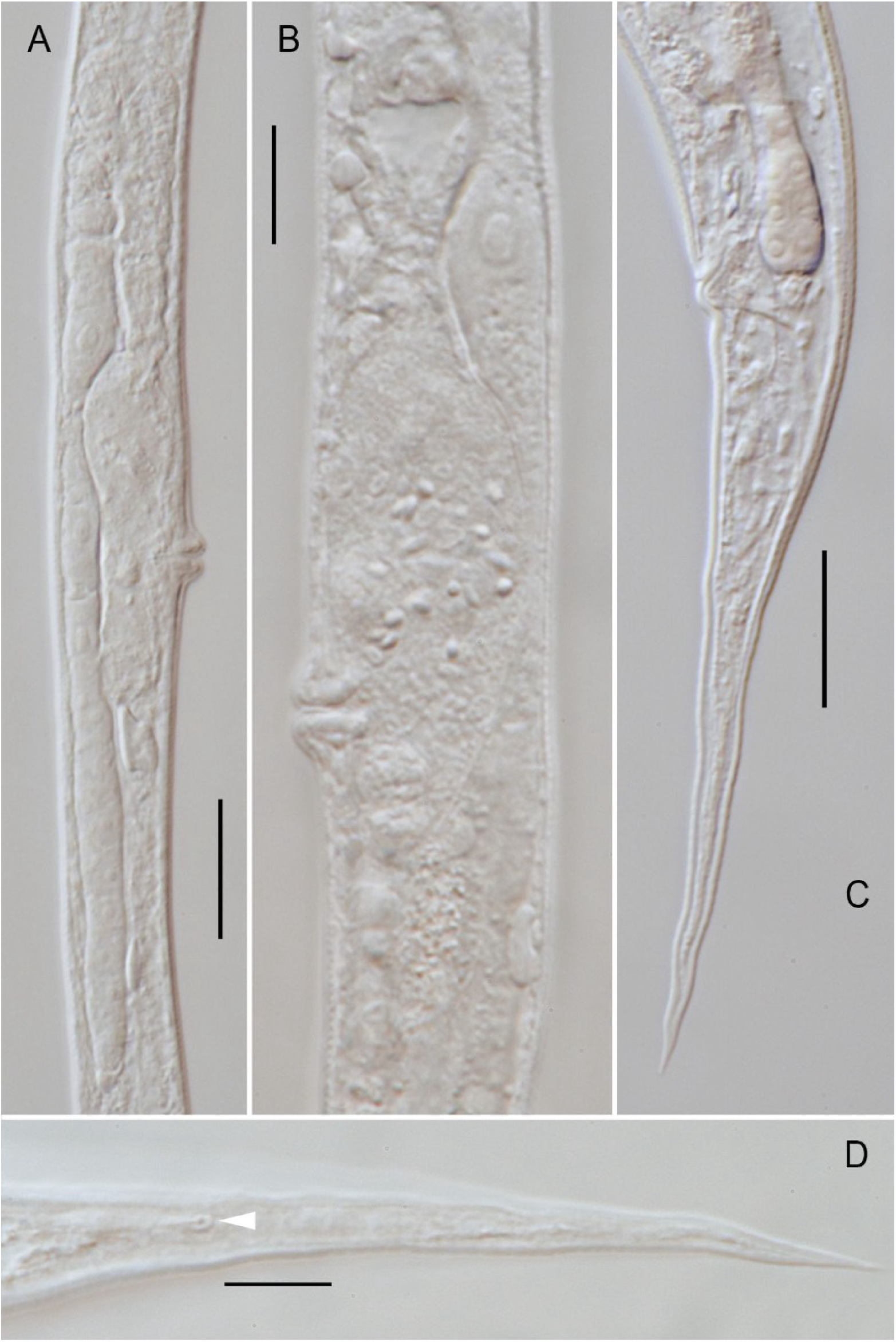
*Neocephalobus halophilus* Paetzold, 1958 (strain BSS8): A. Entire female reproductive system. B: Part of the female reproductive system showing uterus, vulva and post-vulval uterine sac. C: Female tail. D: Phasmid (arrow). Scale bars: A-D = 20 *µ*m.

**Figure 10:**
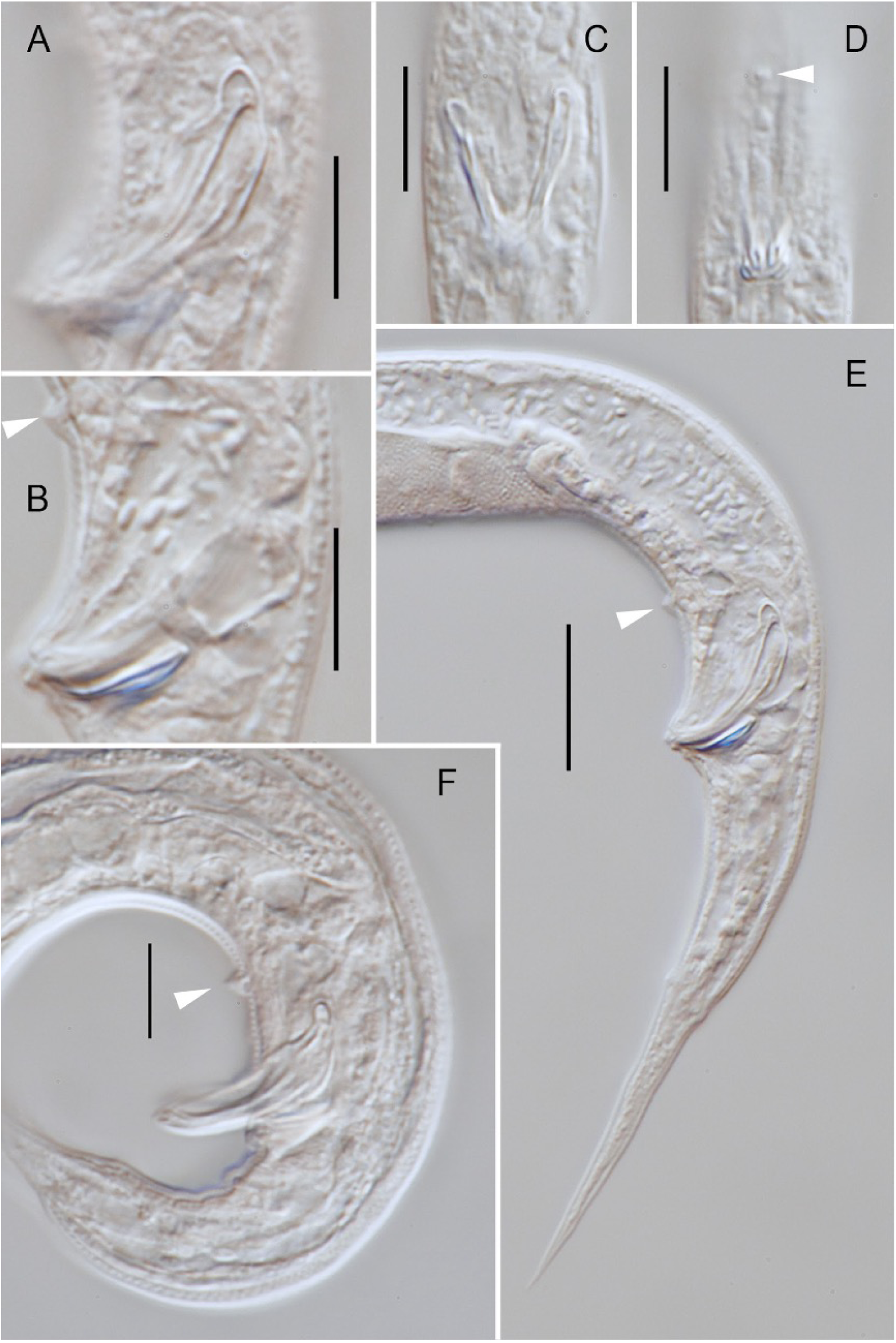
*Neocephalobus halophilus* Paetzold, 1958 (strain BSS8). Lateral (A-B) and ventral (C-D) views of cloacal region. E-F: Lateral view of caudal region. Arrow points to midventral precloacal papilliform sensillum in B and D-F. Scale bars: A-D = 10*µ*m, E-F = 20 *µ*m.

**Figure 11:**
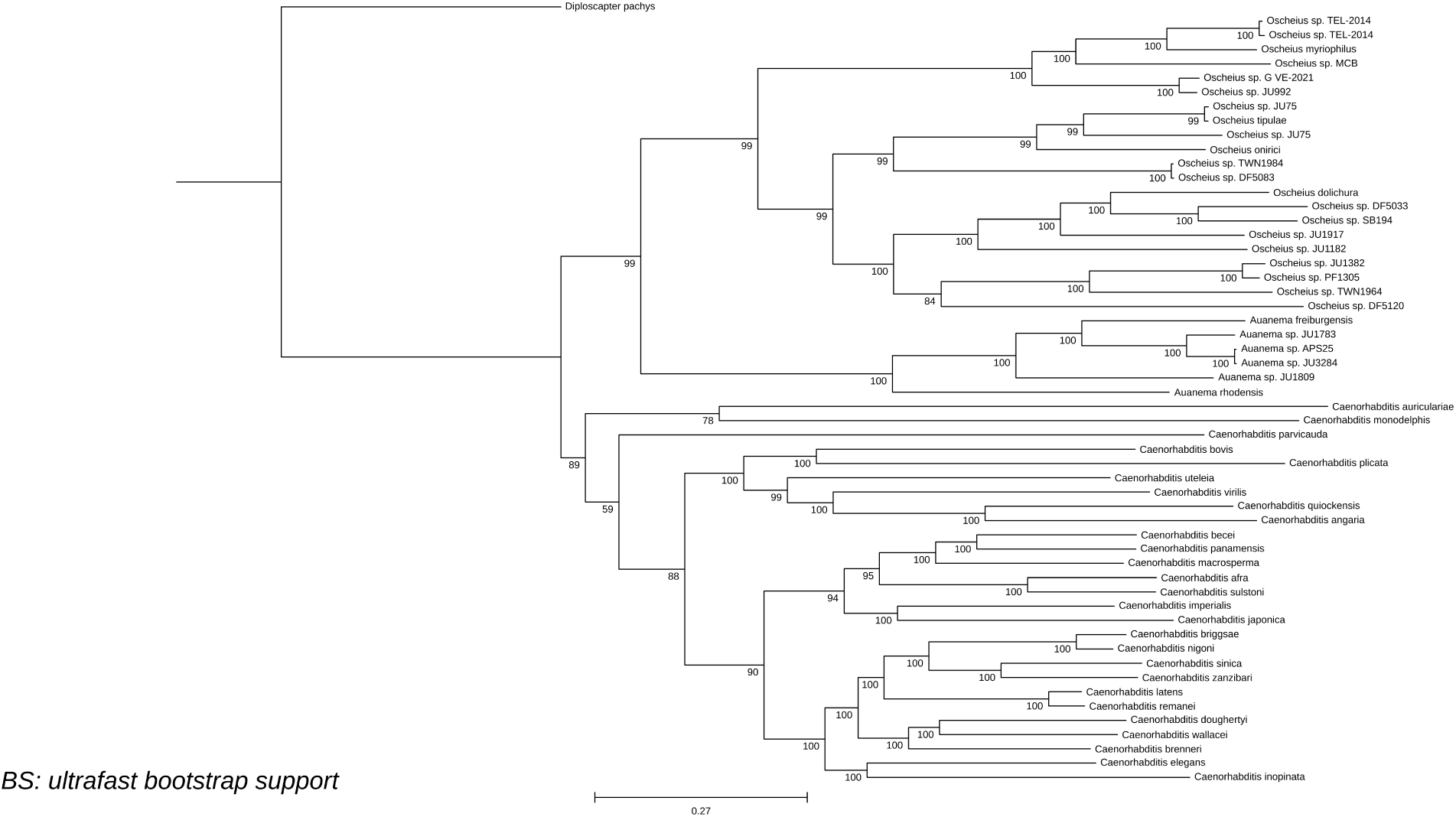
Phylogenetic reconstruction of the Rhabditidae family based on UCEs. *Diploscapter coronatus* is excluded due to very long branch (possibly LBA) and incorrect placing given the low amount of UCEs harvested in the genome. This reconstrucction is based in a 65% occupancy matrix based on 215 alignments.

**Table 7:**
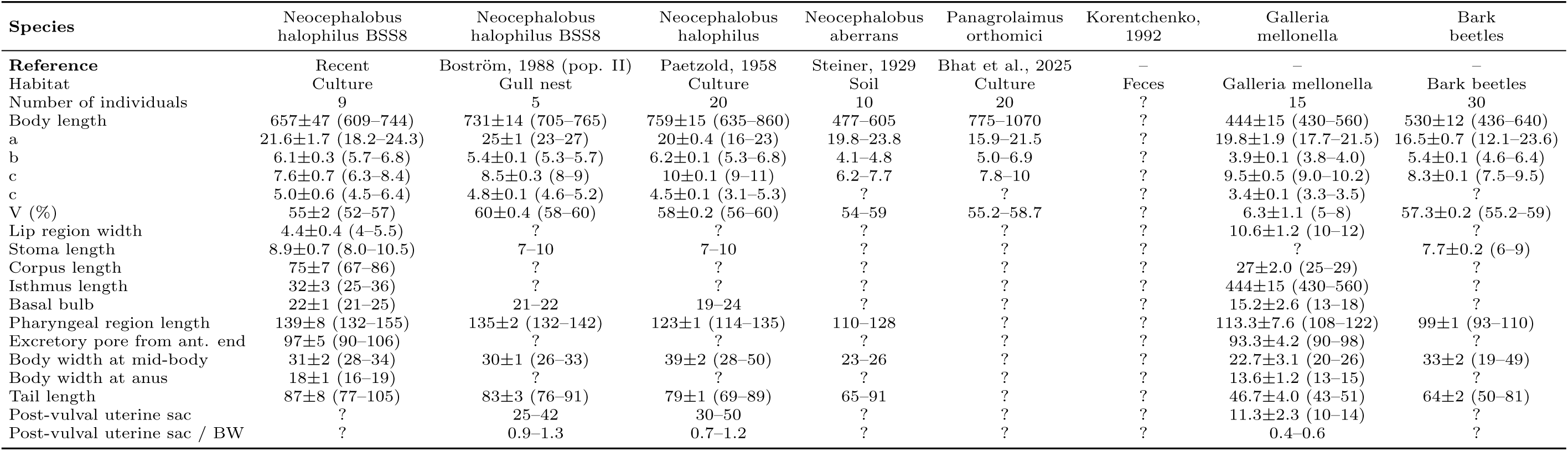
Supplementary Table 7: Morphometrics of *Neocephalobus halophilus* Paetzold, 1958 (strain BSS8), and closely related species (females). Values are means ± standard deviation followed by ranges in parentheses. “?” denotes unavailable data.

**Table 8:**
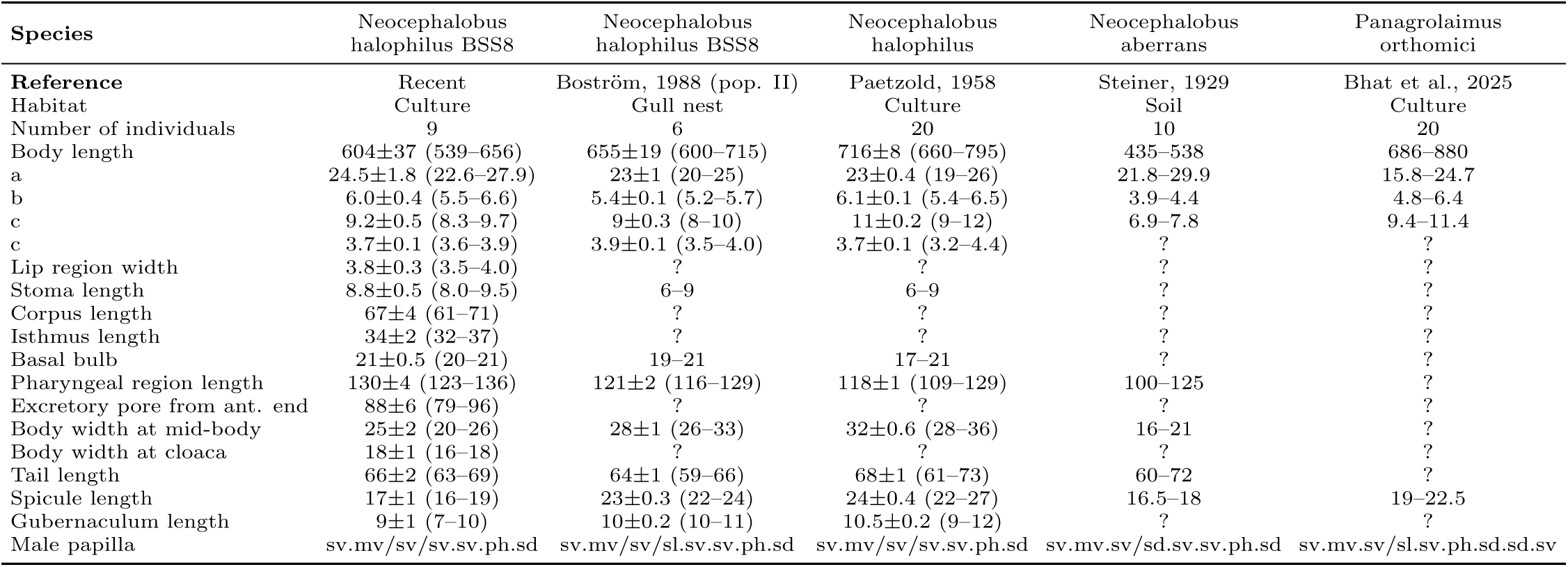
Supplementary Table 8: Morphometrics of *Neocephalobus halophilus* Paetzold, 1958 (strain BSS8), and closely related species (males). Values are means ± standard deviation followed by ranges in parentheses. “?” denotes unavailable data.

**Table 9:** *Oscheius* GenBank accession numbers — Rhabditidae family.

| **Genera** | **GenBank Accession** |
| --- | --- |
| *Oscheius* | GCA_932521025.1 |
| *Oscheius* | GCA_932521405.1 |
| *Oscheius* | GCA_932521415.1 |
| *Oscheius* | GCA_932521035.1 |
| *Oscheius* | GCA_964036215.1 |
| *Oscheius* | GCA_964057215.1 |
| *Oscheius* | GCA_037178975.1 |
| *Oscheius* | GCA_037178855.1 |
| *Oscheius* | GCA_022343465.1 |
| *Oscheius* | GCA_000934875.1 |
| *Oscheius* | GCA_001513535.1 |
| *Oscheius* | GCA_013425905.1 |
| *Oscheius* | GCA_964264085.1 |
| *Oscheius* | GCA_964264175.1 |
| *Oscheius* | GCA_964264275.1 |
| *Oscheius* | GCA_964264245.1 |
| *Oscheius* | GCA_964261225.1 |
| *Oscheius* | GCA_964264255.1 |
| *Oscheius* | GCA_964261325.1 |
| *Oscheius* | GCA_964261265.1 |
| *Oscheius* | GCA_964264185.1 |
| *Oscheius* | GCA_037179065.1 |
| *Oscheius* | GCA_037178915.1 |
| *Oscheius* | GCA_037179185.1 |
| *Oscheius* | GCA_037178995.1 |
| *Oscheius* | GCA_037179095.1 |
| *Oscheius* | GCA_037178805.1 |
| *Oscheius* | GCA_037178875.1 |
| *Oscheius* | GCA_037179085.1 |
| *Oscheius* | GCA_037179045.1 |
| *Oscheius* | GCA_037179165.1 |
| *Oscheius* | GCA_037178765.1 |
| *Oscheius* | GCA_037178905.1 |
| *Oscheius* | GCA_037178795.1 |
| *Oscheius* | GCA_037178965.1 |
| *Oscheius* | GCA_037178985.1 |
| *Oscheius* | GCA_037178755.1 |
| *Oscheius* | GCA_037179145.1 |
| *Oscheius* | GCA_037178945.1 |
| *Oscheius* | GCA_037179075.1 |
| *Oscheius* | GCA_900184235.1 |
| *Oscheius* | GCA_022343475.1 |
| *Oscheius* | GCA_022343505.1 |
| *Oscheius* | GCA_964036205.1 |
| *Oscheius* | GCA_964264195.1 |
| *Oscheius* | GCA_964263735.1 |
| *Oscheius* | GCA_964263365.1 |
| *Oscheius* | GCA_964057235.1 |
| *Oscheius* | GCA_964263805.1 |
| *Oscheius* | GCA_001630785.1 |
| *Oscheius* | GCA_964263935.1 |
| *Oscheius* | GCA_964264165.1 |

**Table 10:**
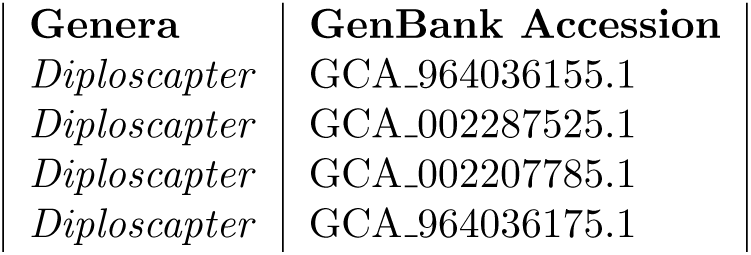
*Diploscapter* GenBank accession numbers – Rhabditidae family.

**Table 11:**
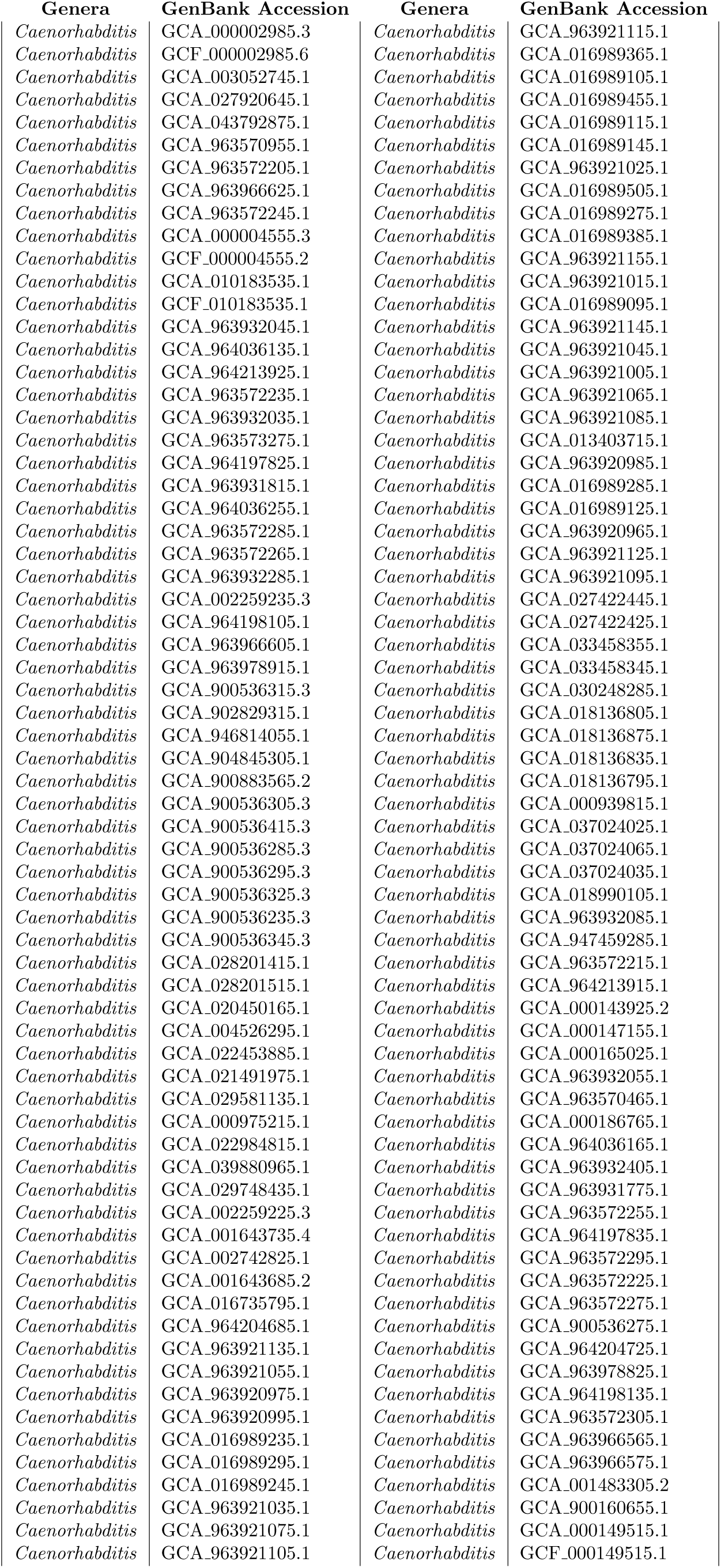
*Caenorhabditis* GenBank accession numbers – Rhabditidae family.

| Genera | GenBank Accession | Genera | GenBank Accession |
| --- | --- | --- | --- |
| *Caenorhabditis* | GCA_000002985.3 | *Caenorhabditis* | GCA_963921115.1 |
| *Caenorhabditis* | GCF_000002985.6 | *Caenorhabditis* | GCA_016989365.1 |
| *Caenorhabditis* | GCA_003052745.1 | *Caenorhabditis* | GCA_016989105.1 |
| *Caenorhabditis* | GCA_027920645.1 | *Caenorhabditis* | GCA_016989455.1 |
| *Caenorhabditis* | GCA_043792875.1 | *Caenorhabditis* | GCA_016989115.1 |
| *Caenorhabditis* | GCA_963570955.1 | *Caenorhabditis* | GCA_016989145.1 |
| *Caenorhabditis* | GCA_963572205.1 | *Caenorhabditis* | GCA_963921025.1 |
| *Caenorhabditis* | GCA_963966625.1 | *Caenorhabditis* | GCA_016989505.1 |
| *Caenorhabditis* | GCA_963572245.1 | *Caenorhabditis* | GCA_016989275.1 |
| *Caenorhabditis* | GCA_000004555.3 | *Caenorhabditis* | GCA_016989385.1 |
| *Caenorhabditis* | GCF_000004555.2 | *Caenorhabditis* | GCA_963921155.1 |
| *Caenorhabditis* | GCA_010183535.1 | *Caenorhabditis* | GCA_963921015.1 |
| *Caenorhabditis* | GCF_010183535.1 | *Caenorhabditis* | GCA_016989095.1 |
| *Caenorhabditis* | GCA_963932045.1 | *Caenorhabditis* | GCA_963921145.1 |
| *Caenorhabditis* | GCA_964036135.1 | *Caenorhabditis* | GCA_963921045.1 |
| *Caenorhabditis* | GCA_964213925.1 | *Caenorhabditis* | GCA_963921005.1 |
| *Caenorhabditis* | GCA_963572235.1 | *Caenorhabditis* | GCA_963921065.1 |
| *Caenorhabditis* | GCA_963932035.1 | *Caenorhabditis* | GCA_963921085.1 |
| *Caenorhabditis* | GCA_963573275.1 | *Caenorhabditis* | GCA_013403715.1 |
| *Caenorhabditis* | GCA_964197825.1 | *Caenorhabditis* | GCA_963920985.1 |
| *Caenorhabditis* | GCA_963931815.1 | *Caenorhabditis* | GCA_016989285.1 |
| *Caenorhabditis* | GCA_964036255.1 | *Caenorhabditis* | GCA_016989125.1 |
| *Caenorhabditis* | GCA_963572285.1 | *Caenorhabditis* | GCA_963920965.1 |
| *Caenorhabditis* | GCA_963572265.1 | *Caenorhabditis* | GCA_963921125.1 |
| *Caenorhabditis* | GCA_963932285.1 | *Caenorhabditis* | GCA_963921095.1 |
| *Caenorhabditis* | GCA_002259235.3 | *Caenorhabditis* | GCA_027422445.1 |
| *Caenorhabditis* | GCA_964198105.1 | *Caenorhabditis* | GCA_027422425.1 |
| *Caenorhabditis* | GCA_963966605.1 | *Caenorhabditis* | GCA_033458355.1 |
| *Caenorhabditis* | GCA_963978915.1 | *Caenorhabditis* | GCA_033458345.1 |
| *Caenorhabditis* | GCA_900536315.3 | *Caenorhabditis* | GCA_030248285.1 |
| *Caenorhabditis* | GCA_902829315.1 | *Caenorhabditis* | GCA_018136805.1 |
| *Caenorhabditis* | GCA_946814055.1 | *Caenorhabditis* | GCA_018136875.1 |
| *Caenorhabditis* | GCA_904845305.1 | *Caenorhabditis* | GCA_018136835.1 |
| *Caenorhabditis* | GCA_900883565.2 | *Caenorhabditis* | GCA_018136795.1 |
| *Caenorhabditis* | GCA_900536305.3 | *Caenorhabditis* | GCA_000939815.1 |
| *Caenorhabditis* | GCA_900536415.3 | *Caenorhabditis* | GCA_037024025.1 |
| *Caenorhabditis* | GCA_900536285.3 | *Caenorhabditis* | GCA_037024065.1 |
| *Caenorhabditis* | GCA_900536295.3 | *Caenorhabditis* | GCA_037024035.1 |
| *Caenorhabditis* | GCA_900536325.3 | *Caenorhabditis* | GCA_018990105.1 |
| *Caenorhabditis* | GCA_900536235.3 | *Caenorhabditis* | GCA_963932085.1 |
| *Caenorhabditis* | GCA_900536345.3 | *Caenorhabditis* | GCA_947459285.1 |
| *Caenorhabditis* | GCA_028201415.1 | *Caenorhabditis* | GCA_963572215.1 |
| *Caenorhabditis* | GCA_028201515.1 | *Caenorhabditis* | GCA_964213915.1 |
| *Caenorhabditis* | GCA_020450165.1 | *Caenorhabditis* | GCA_000143925.2 |
| *Caenorhabditis* | GCA_004526295.1 | *Caenorhabditis* | GCA_000147155.1 |
| *Caenorhabditis* | GCA_022453885.1 | *Caenorhabditis* | GCA_000165025.1 |
| *Caenorhabditis* | GCA_021491975.1 | *Caenorhabditis* | GCA_963932055.1 |
| *Caenorhabditis* | GCA_029581135.1 | *Caenorhabditis* | GCA_963570465.1 |
| *Caenorhabditis* | GCA_000975215.1 | *Caenorhabditis* | GCA_000186765.1 |
| *Caenorhabditis* | GCA_022984815.1 | *Caenorhabditis* | GCA_964036165.1 |
| *Caenorhabditis* | GCA_039880965.1 | *Caenorhabditis* | GCA_963932405.1 |
| *Caenorhabditis* | GCA_029748435.1 | *Caenorhabditis* | GCA_963931775.1 |
| *Caenorhabditis* | GCA_002259225.3 | *Caenorhabditis* | GCA_963572255.1 |
| *Caenorhabditis* | GCA_001643735.4 | *Caenorhabditis* | GCA_964197835.1 |
| *Caenorhabditis* | GCA_002742825.1 | *Caenorhabditis* | GCA_963572295.1 |
| *Caenorhabditis* | GCA_001643685.2 | *Caenorhabditis* | GCA_963572225.1 |
| *Caenorhabditis* | GCA_016735795.1 | *Caenorhabditis* | GCA_963572275.1 |
| *Caenorhabditis* | GCA_964204685.1 | *Caenorhabditis* | GCA_900536275.1 |
| *Caenorhabditis* | GCA_963921135.1 | *Caenorhabditis* | GCA_964204725.1 |
| *Caenorhabditis* | GCA_963921055.1 | *Caenorhabditis* | GCA_963978825.1 |
| *Caenorhabditis* | GCA_963920975.1 | *Caenorhabditis* | GCA_964198135.1 |
| *Caenorhabditis* | GCA_963920995.1 | *Caenorhabditis* | GCA_963572305.1 |
| *Caenorhabditis* | GCA_016989235.1 | *Caenorhabditis* | GCA_963966565.1 |
| *Caenorhabditis* | GCA_016989295.1 | *Caenorhabditis* | GCA_963966575.1 |
| *Caenorhabditis* | GCA_016989245.1 | *Caenorhabditis* | GCA_001483305.2 |
| *Caenorhabditis* | GCA_963921035.1 | *Caenorhabditis* | GCA_900160655.1 |
| *Caenorhabditis* | GCA_963921075.1 | *Caenorhabditis* | GCA_000149515.1 |
| *Caenorhabditis* | GCA_963921105.1 | *Caenorhabditis* | GCF_000149515.1 |

**Table 12:** *Auanema* GenBank accession numbers – Rhabditidae family.

| **Genera** | **GenBank Accession** |
| --- | --- |
| *Auanema* | GCA_964057225.1 |
| *Auanema* | GCA_030370435.1 |
| *Auanema* | GCA_943334845.2 |
| *Auanema* | GCA_964264295.1 |
| *Auanema* | GCA_964263715.1 |
| *Auanema* | GCA_964263695.1 |
| *Auanema* | GCA_947366455.1 |
| *Auanema* | GCA_964057245.1 |
| *Auanema* | GCA_964263765.1 |

#### A.3 Distribution and characteristics of UCEs in Rhabditidae and Panagrolaimidae

We assessed the distribution of Ultra-Conserved Elements (UCEs) across strains in the families Rhabditidae and Panagrolaimidae by analyzing the total number of UCEs per strain. The results, presented in Figures 13 and 12, reveal considerable variation in UCE counts among strains within each family. Specifically, in Rhabditidae, UCE counts range from 1 to 5700, with a median of 3840, while in Panagrolaimidae, they range from 15 to 1457, with a median of 790.0.

**Figure 12:**
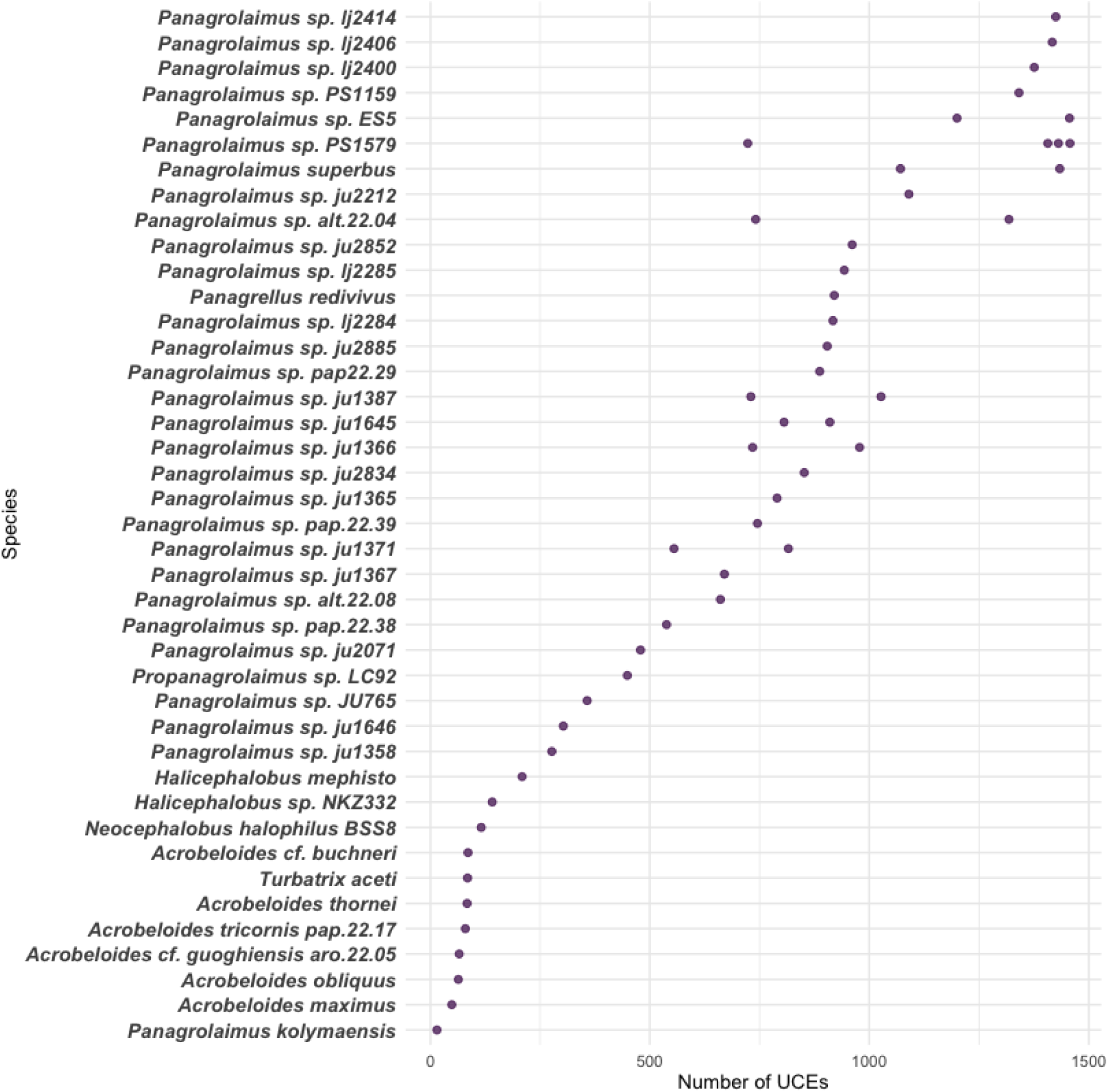
Number of UCEs per species in Panagrolaimidae. Species are sorted from lowest to highest UCE count.

**Figure 13:**
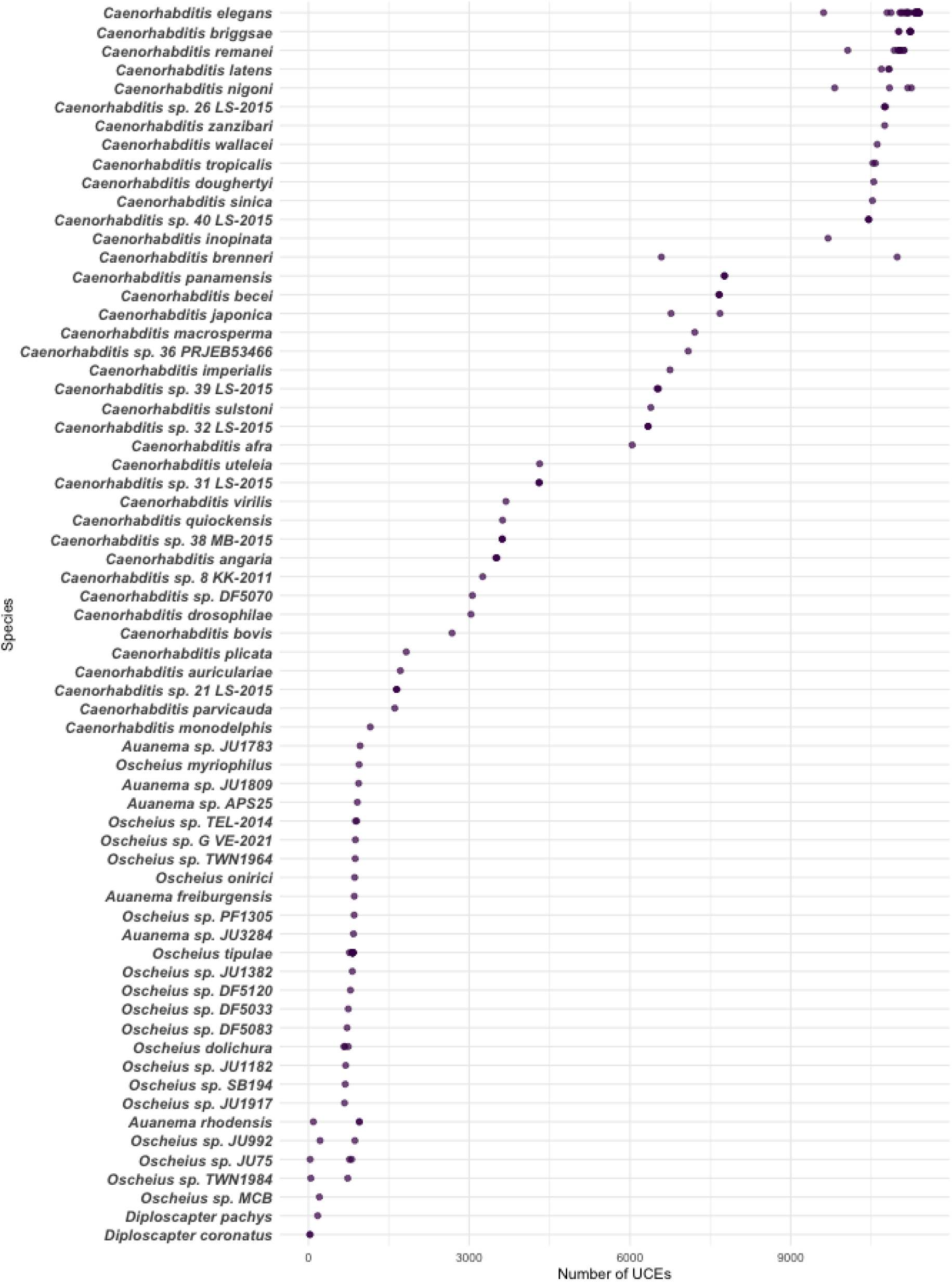
Number of UCEs per species in Rhabditidae. Species are sorted from lowest to highest UCE count.

##### A.3.1 UCEs shared across all the genera in the Rhabditidae and Panagrolaimidae family

**Table 13:** Shared UCEs across all genera for each family.

| Family | Total number | UCE name |
| --- | --- | --- |
| Rhabditidae | 50 | uce.1269, uce.16032, uce.1524, uce.1260, uce.1673, uce.1892, uce.14291, uce.15873, uce.512, uce.16435, uce.19283, uce.7014, uce.15882, uce.262, uce.1190, uce.1271, uce.13410, uce.655, uce.1272, uce.14817, uce.424, uce.5519, uce.2159, uce.2212, uce.2407, uce.583, uce.1309, uce.4011, uce.16149, uce.6068, uce.2630, uce.731, uce.16271, uce.17033, uce.1326, uce.16433, uce.2401, uce.3013, uce.7820, uce.739, uce.1319, uce.15494, uce.18743, uce.2484, uce.15826, uce.737, uce.1739, uce.1905, uce.1984, uce.18176 |
| Panagrolaimidae | 84 | uce.2078, uce.1795, uce.5994, uce.5306, uce.8657, uce.8674, uce.8695, uce.8786, uce.9007, uce.10095, uce.10105, uce.10196, uce.8702, uce.10397, uce.281, uce.284, uce.431, uce.459, uce.604, uce.618, uce.663, uce.9794, uce.7942, uce.4462, uce.6371, uce.6493, uce.7157, uce.5355, uce.5498, uce.5515, uce.4246, uce.9888, uce.9910, uce.4359, uce.4261, uce.10506, uce.10578, uce.7322, uce.7567, uce.7646, uce.7753, uce.7765, uce.6749, uce.10627, uce.6806, uce.6606, uce.7323, uce.8091, uce.8113, uce.8114, uce.8234, uce.4024, uce.4162, uce.5069, uce.8318, uce.8373, uce.8398, uce.8429, uce.9348, uce.9422, uce.9427, uce.4491, uce.4493, uce.4559, uce.4655, uce.5655, uce.5769, uce.5836, uce.5845, uce.9316, uce.4329, uce.6370, uce.7084, uce.323, uce.5837, uce.7309, uce.7996, uce.3, uce.194, uce.1316, uce.10496, uce.227, uce.6826, uce.5043 |

#### A.4 Benchmark of Machine Learning Models on Rhabditidae Data

A comprehensive comparison of the models is provided through an AUC comparison plot, illustrating the relative performance of each approach, see Figure 14.

**Figure 14:**
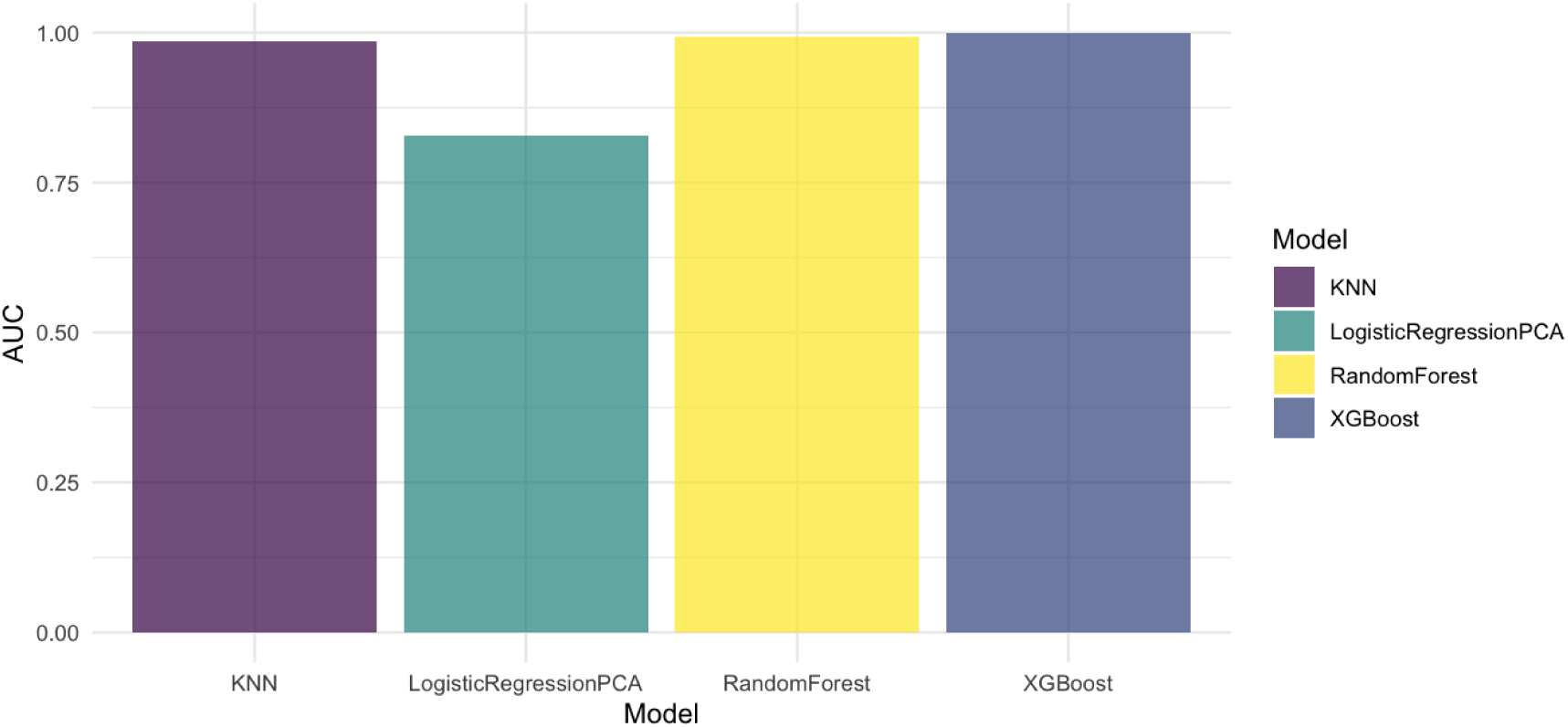
Comparison of AUC scores across ML models on Rhabditidae data.

Complete class-specific performance metrics are included, detailing sensitivity, specificity, precision, recall, and balanced accuracy. Additionally, confusion matrix heatmaps for each model offer visual insights into classification accuracy across all genera, see Figure 15.

##### Random Forest

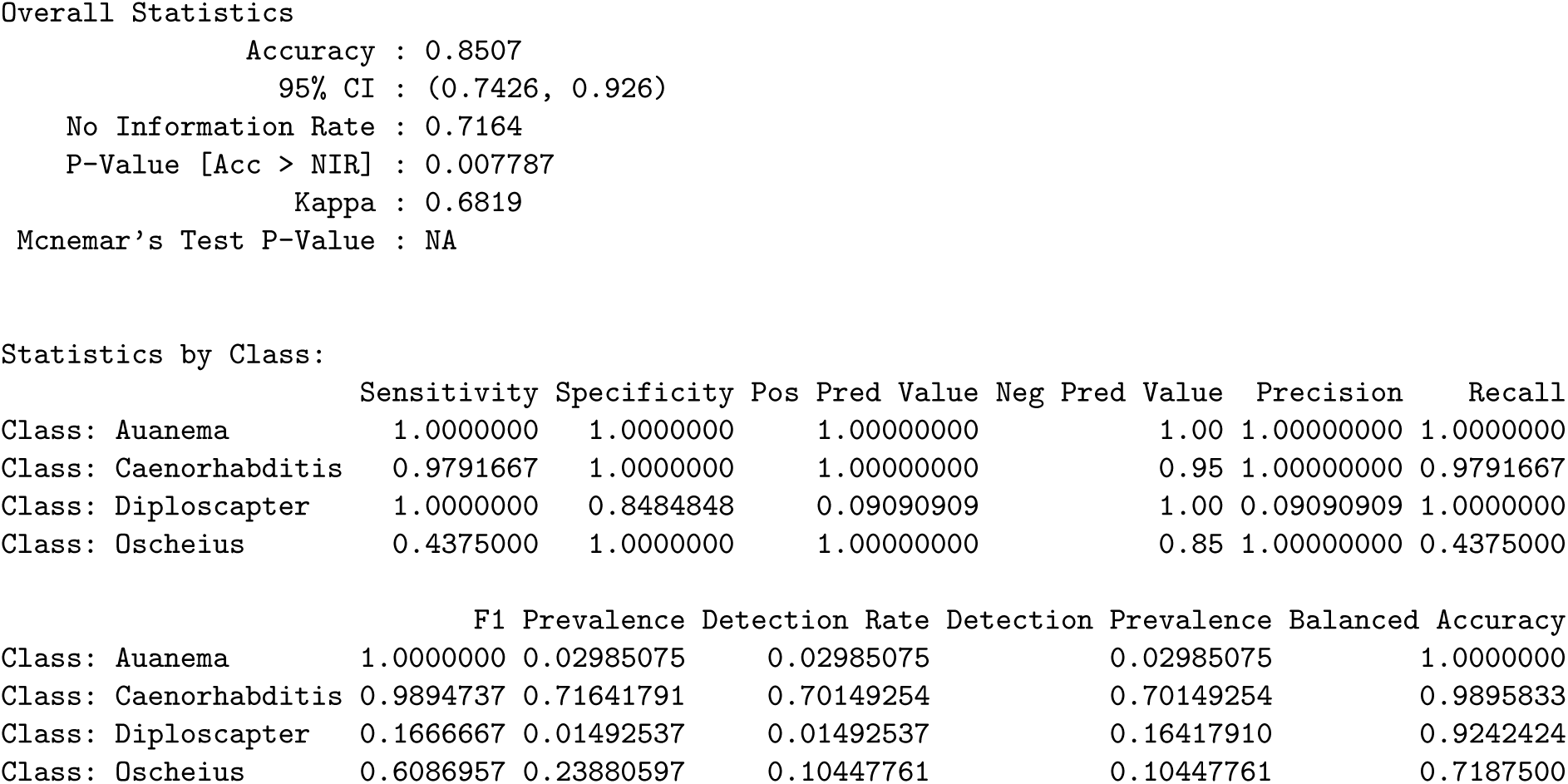

##### Logistic Regression with PCA

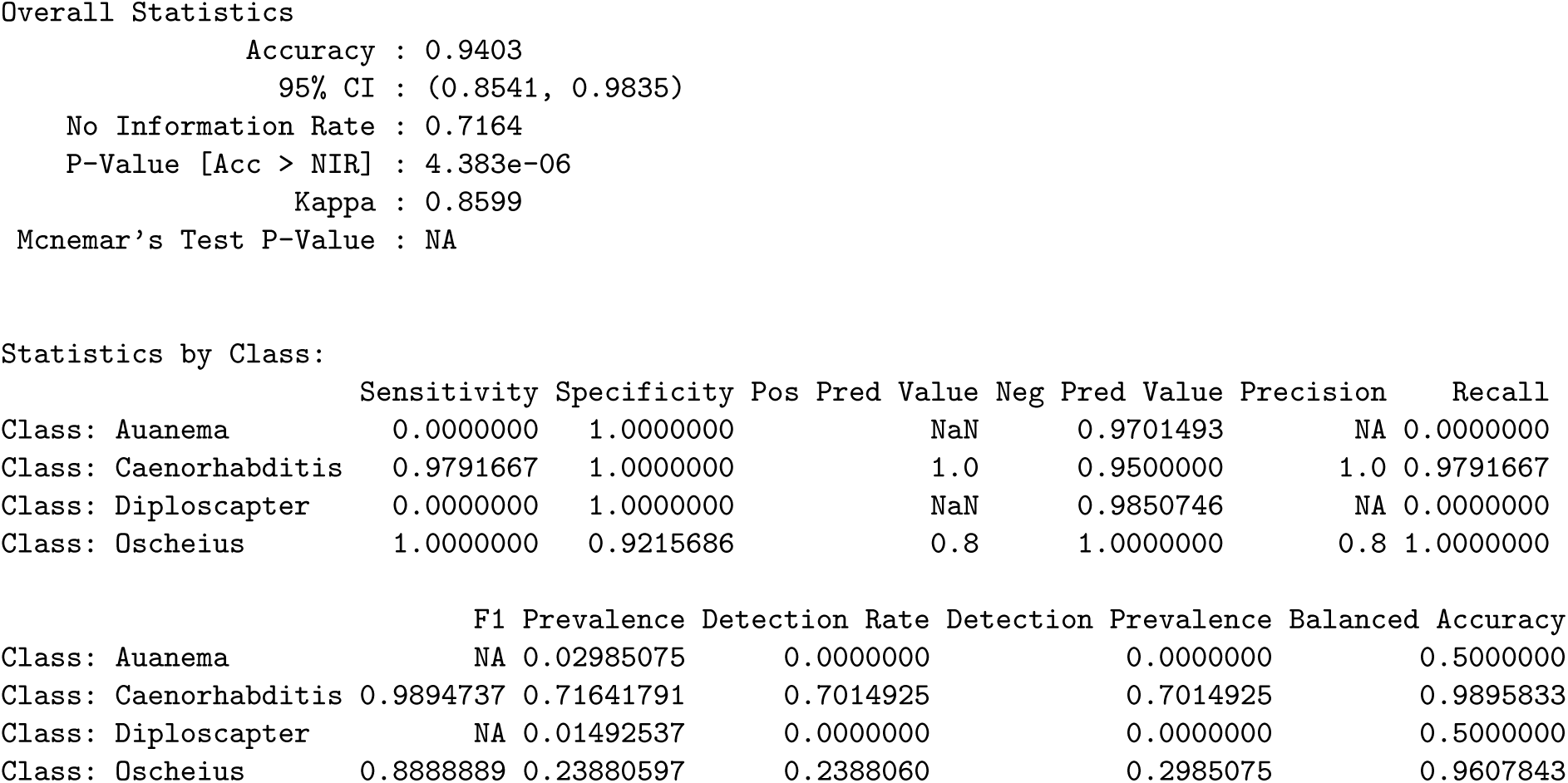

##### k-Nearest Neighbors

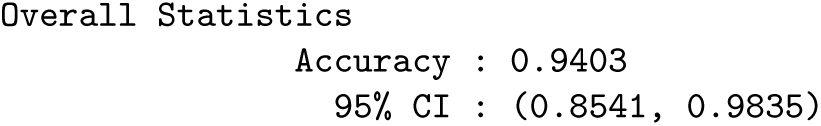
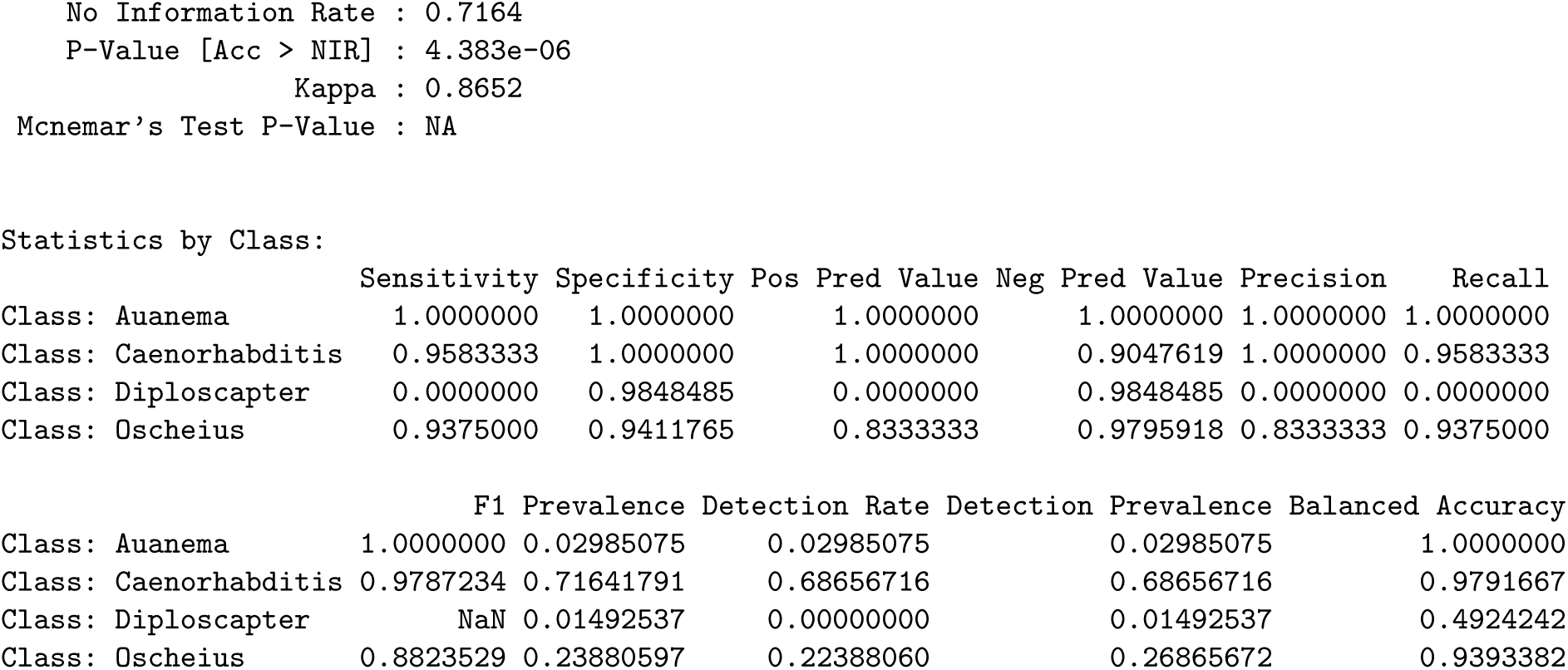

##### XGBoost

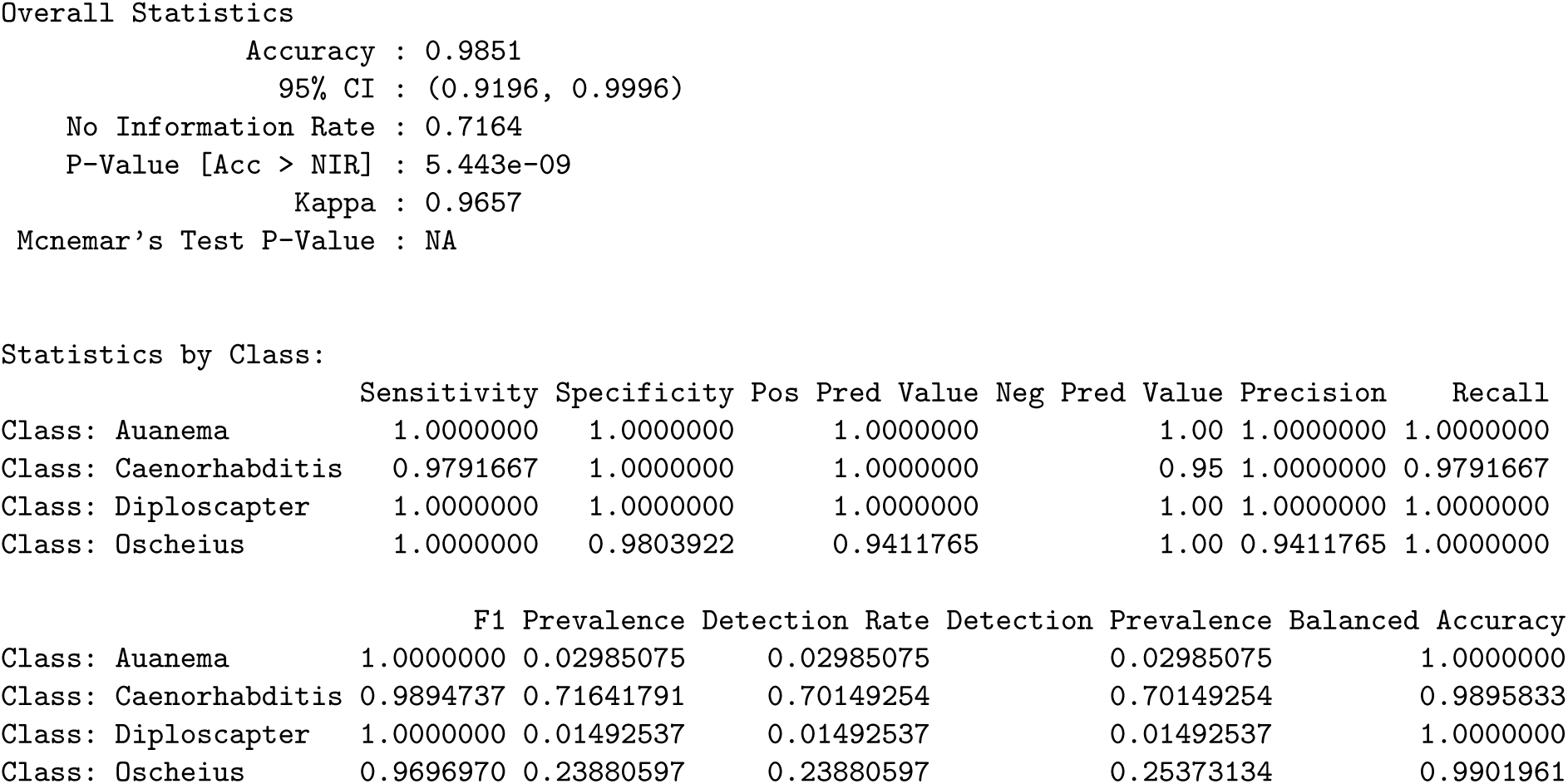

##### A.4.1 Identification of important features using non-zero importance scores in XGBoost for Rhabditidae family

A cross-validation procedure was conducted to assess the impact of feature count on model performance. The process involved iteratively training the XGBoost model with increasing numbers of top-ranked features, as determined by their importance scores. Specifically, the model was trained and evaluated using feature subsets ranging from the single most important feature up to the top 20 features. For each iteration, the Area Under the Curve (AUC) was calculated using the ‘multiclass.roc‘ function from the ‘pROC’ package, providing a measure of the model’s predictive accuracy. The resulting AUC values were then plotted against the number of features used, generating a visual representation of the relationship between feature count and model performance.

**Figure 15:**
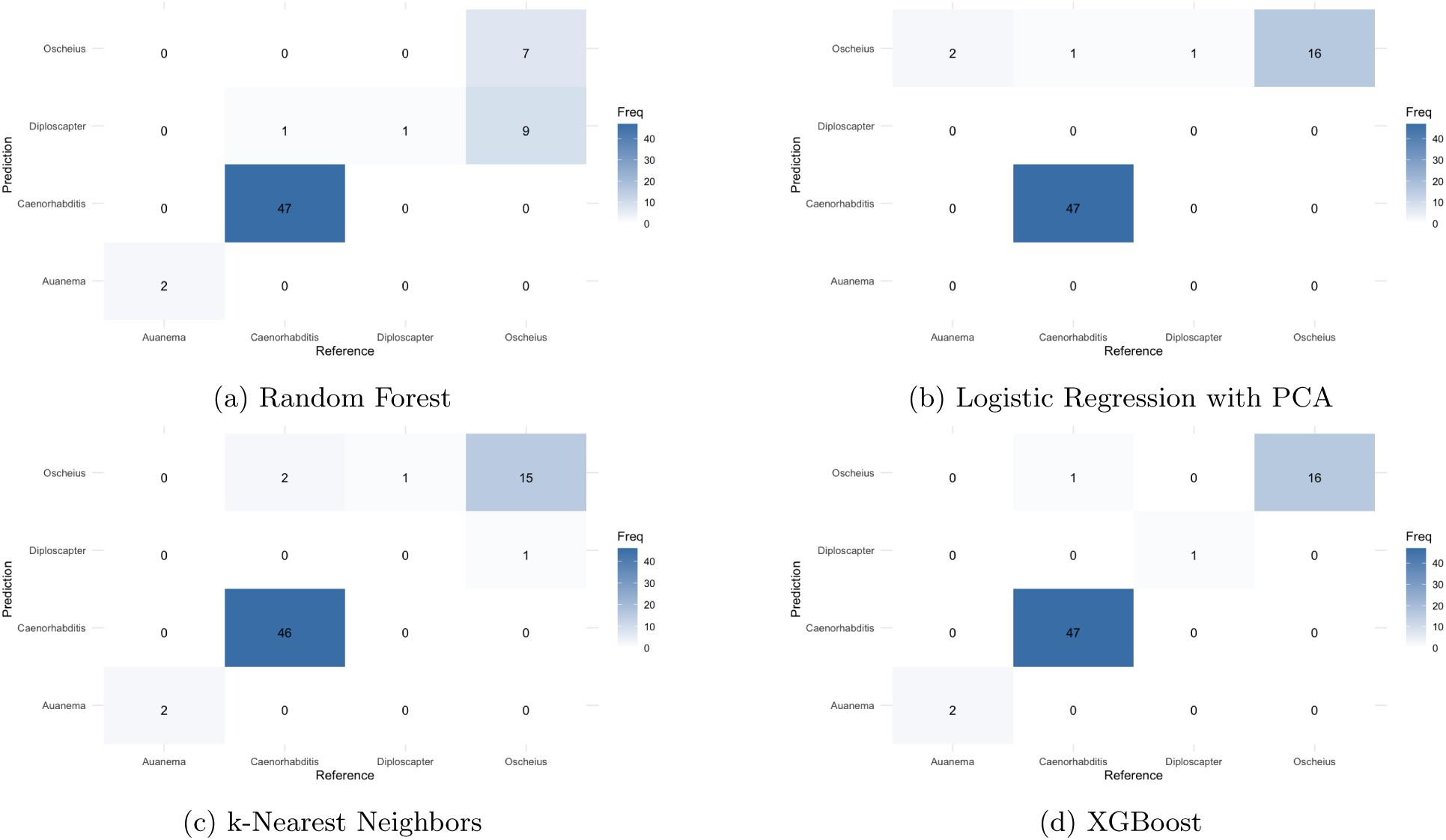
Confusion matrix heatmaps for different ML models on Rhabditidae Data.

**Figure 16:**
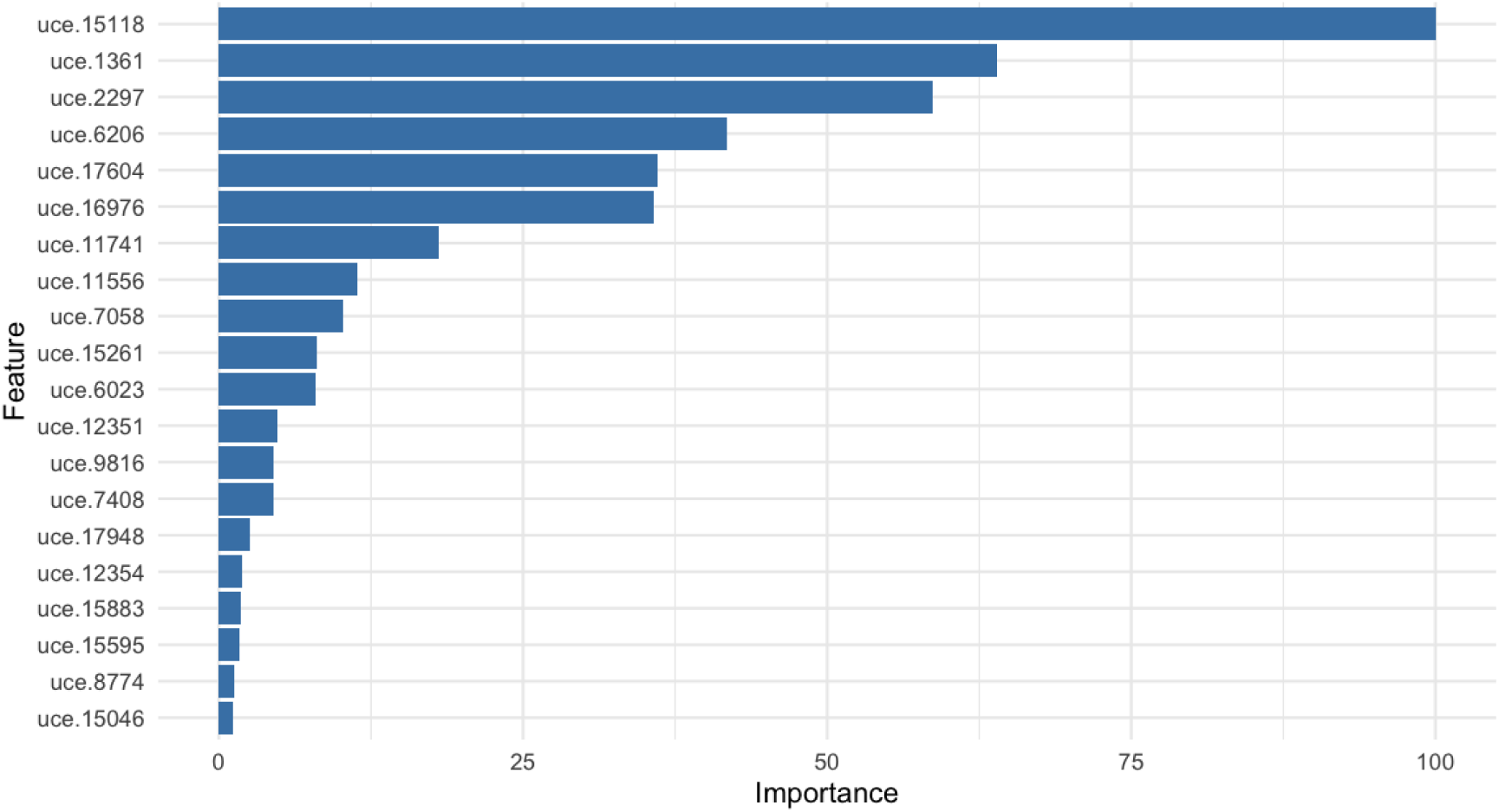
Top 20 features ranked by importance scores (XGBoost) for Rhabditidae family.

**Table 14:** Feature importance scores (XGBoost) on the Rhabditidae family, sorted in descending order.

| # | Feature | Overall | # | Feature | Overall |
| --- | --- | --- | --- | --- | --- |
| 1 | uce.15118 | 100.00000000 | 24 | uce.3117 | 0.57467362 |
| 2 | uce.1361 | 63.98891640 | 25 | uce.1508 | 0.55899331 |
| 3 | uce.2297 | 58.70899293 | 26 | uce.377 | 0.54091615 |
| 4 | uce.6206 | 41.76429892 | 27 | uce.6314 | 0.40157400 |
| 5 | uce.17604 | 36.08823354 | 28 | uce.9054 | 0.29854266 |
| 6 | uce.16976 | 35.71085919 | 29 | uce.7504 | 0.29501397 |
| 7 | uce.11741 | 18.01152015 | 30 | uce.15693 | 0.28641136 |
| 8 | uce.11556 | 11.37746178 | 31 | uce.1739 | 0.26911590 |
| 9 | uce.7058 | 10.21175252 | 32 | uce.4435 | 0.26010081 |
| 10 | uce.15261 | 8.06170688 | 33 | uce.14951 | 0.20215311 |
| 11 | uce.6023 | 7.97789374 | 34 | uce.739 | 0.19555710 |
| 12 | uce.12351 | 4.82092405 | 35 | uce.5314 | 0.17701833 |
| 13 | uce.9816 | 4.52307700 | 36 | uce.16080 | 0.17486536 |
| 14 | uce.7408 | 4.45983387 | 37 | uce.9712 | 0.16037217 |
| 15 | uce.17948 | 2.56738332 | 38 | uce.16977 | 0.14765256 |
| 16 | uce.12354 | 1.90536817 | 39 | uce.12716 | 0.13513669 |
| 17 | uce.15883 | 1.84721198 | 40 | uce.15269 | 0.12088749 |
| 18 | uce.15595 | 1.69534444 | 41 | uce.10818 | 0.11498949 |
| 19 | uce.8774 | 1.26314020 | 42 | uce.13654 | 0.07959777 |
| 20 | uce.15046 | 1.13152980 | 43 | uce.8269 | 0.06693109 |
| 21 | uce.16507 | 1.08547413 | 44 | uce.9042 | 0.04954661 |
| 22 | uce.12197 | 0.96881894 | 45 | uce.16032 | 0.03010273 |
| 23 | uce.14879 | 0.88733092 | 46 | uce.12732 | 0.02833252 |

#### A.5 Panagrolaimidae classification model performance

**Figure 17:**
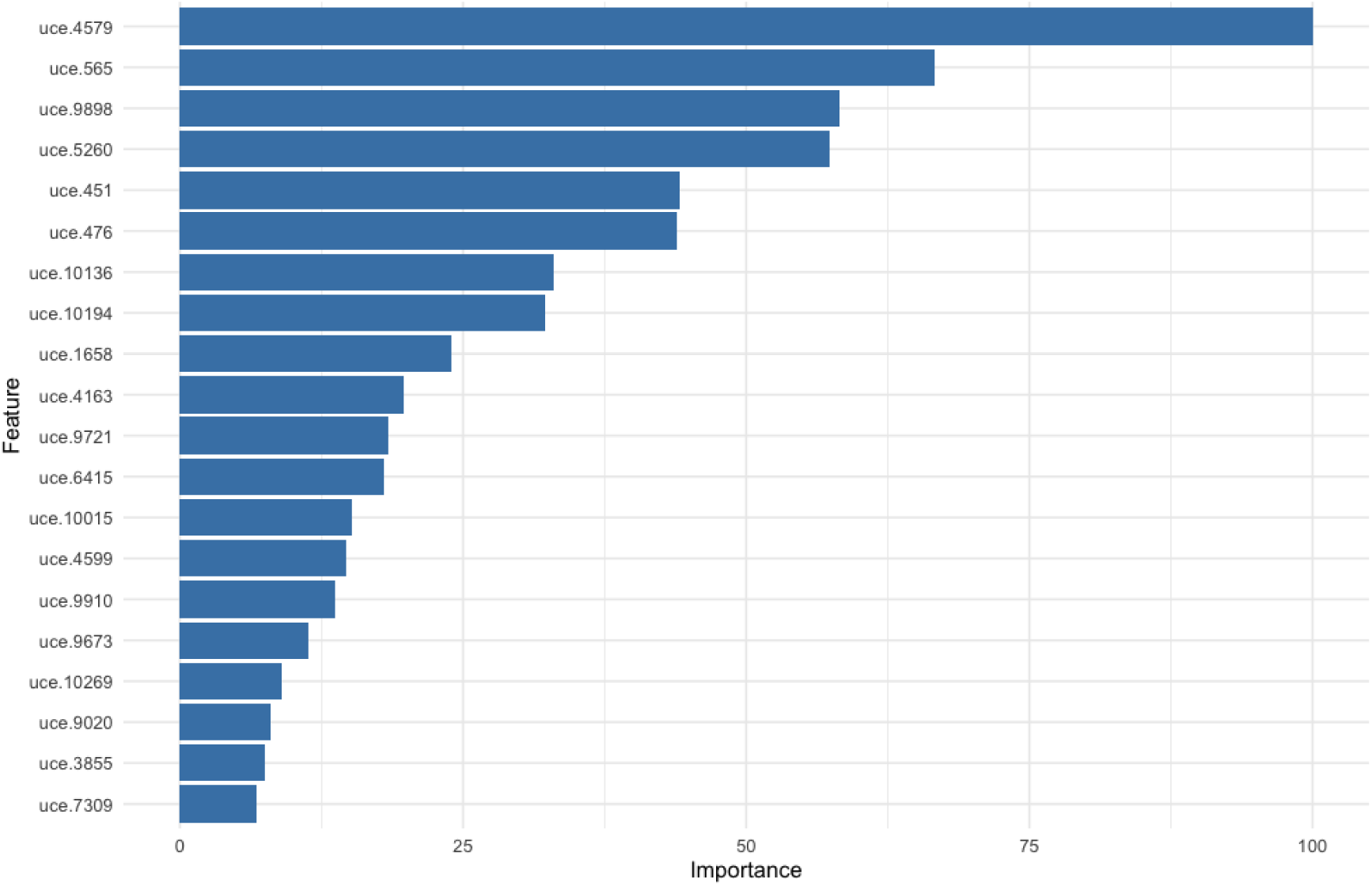
Top 20 features ranked by importance scores (XGBoost) for Panagrolaimidae family.

**Table 15:** Top UCE feature importance and their overall scores for the Panagrolaimidae family.

| # | Feature | Overall | # | Feature | Overall | # | Feature | Overall |
| --- | --- | --- | --- | --- | --- | --- | --- | --- |
| 1 | uce.4579 | 100.000000 | 14 | uce.4599 | 14.631350 | 27 | uce.459 | 5.041798 |
| 2 | uce.565 | 66.623360 | 15 | uce.9910 | 13.704530 | 28 | uce.215 | 3.982949 |
| 3 | uce.9898 | 58.208410 | 16 | uce.9673 | 11.354810 | 29 | uce.7806 | 3.966508 |
| 4 | uce.5260 | 57.338690 | 17 | uce.10269 | 9.037717 | 30 | uce.9218 | 3.224067 |
| 5 | uce.451 | 44.179020 | 18 | uce.9020 | 7.986509 | 31 | uce.4381 | 3.149135 |
| 6 | uce.476 | 43.890580 | 19 | uce.3855 | 7.445594 | 32 | uce.6949 | 2.629612 |
| 7 | uce.10136 | 32.933190 | 20 | uce.7309 | 6.782274 | 33 | uce.8786 | 2.131640 |
| 8 | uce.10194 | 32.206180 | 21 | uce.7825 | 6.602921 | 34 | uce.6827 | 2.059504 |
| 9 | uce.1658 | 23.978340 | 22 | uce.10532 | 6.213068 | 35 | uce.7163 | 2.057336 |
| 10 | uce.4163 | 19.712390 | 23 | uce.194 | 6.027681 | 36 | uce.6376 | 1.980775 |
| 11 | uce.9721 | 18.340240 | 24 | uce.8785 | 5.904270 | 37 | uce.9291 | 1.826193 |
| 12 | uce.6415 | 18.004650 | 25 | uce.5243 | 5.803961 | 38 | uce.4230 | 1.572708 |
| 13 | uce.10015 | 15.175680 | 26 | uce.8674 | 5.418900 | 39 | uce.323 | 1.553780 |

##### XGBoost – full features Panagrolaimidae

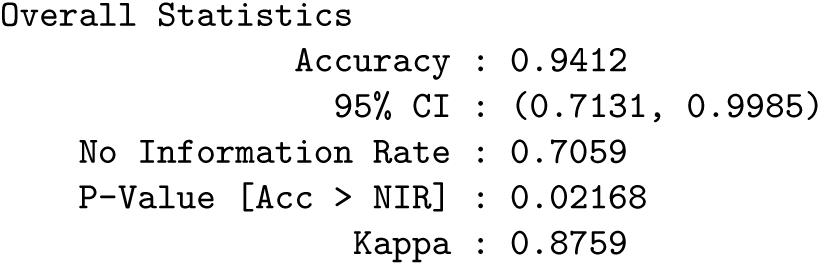
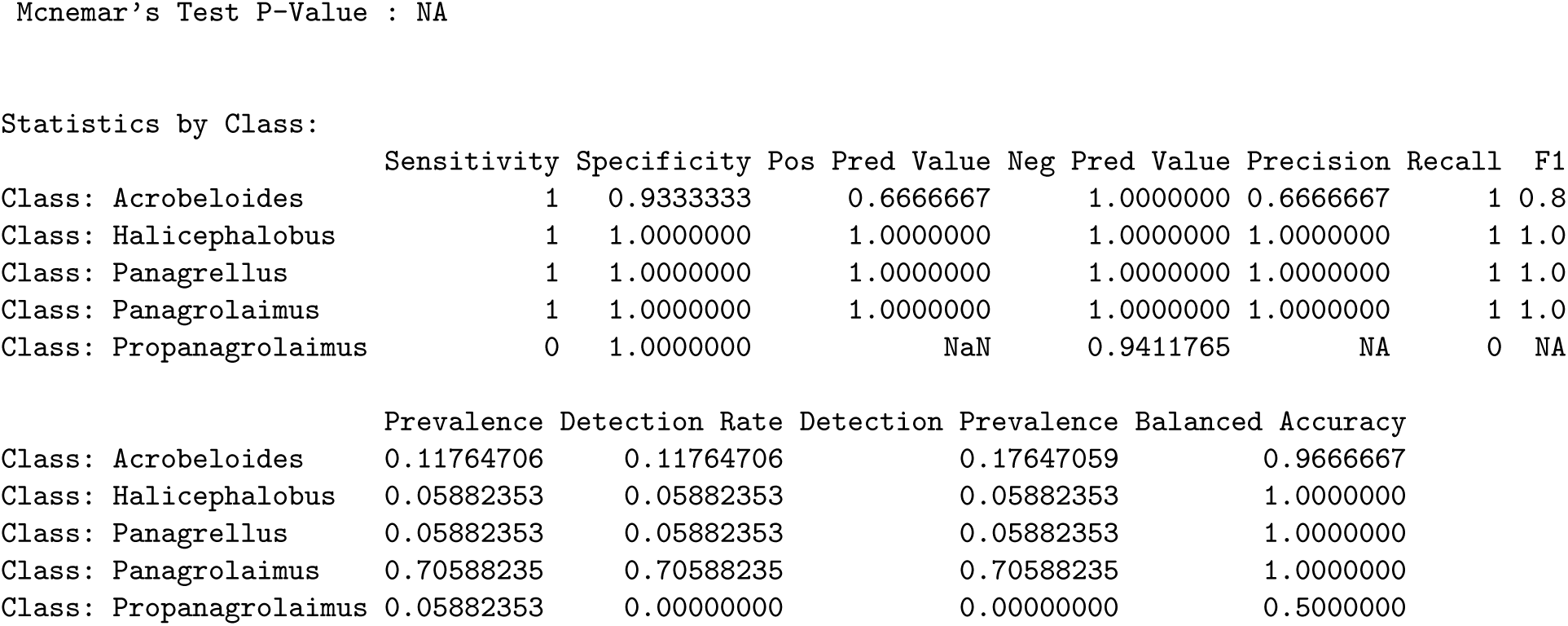

##### XGBoost – feature importance Panagrolaimidae

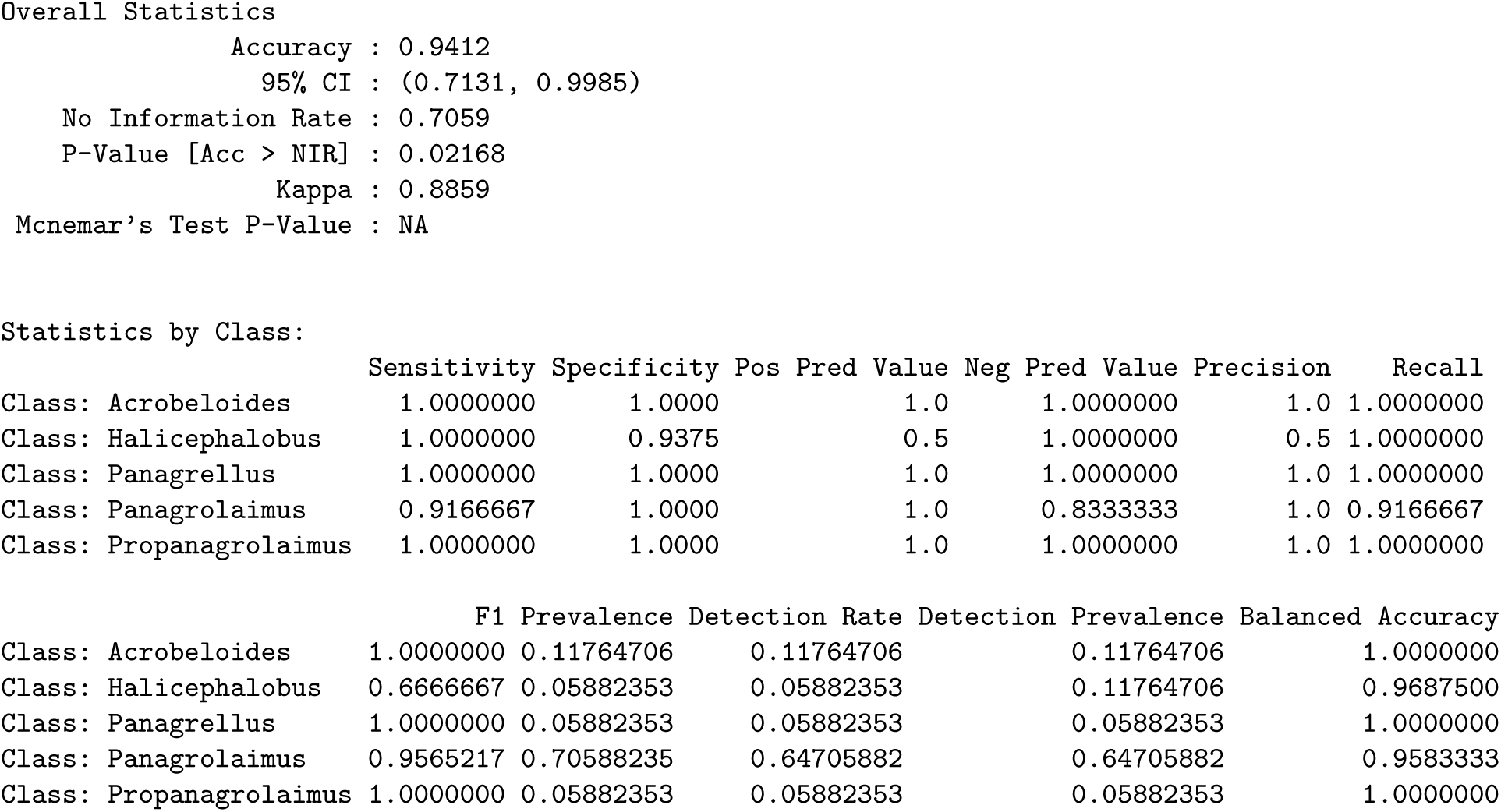

**Figure 18:**
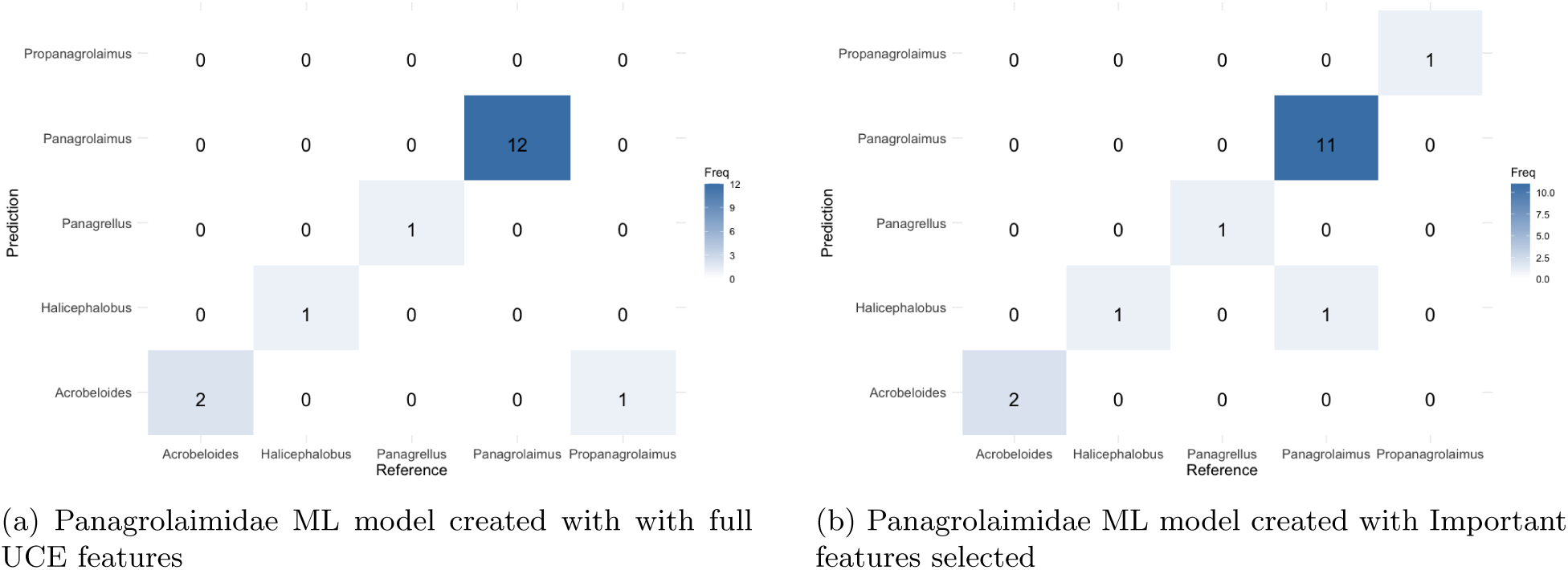
Confusion matrices in Panagrolaimidae.

#### A.6 Analysis of Common UCEs

##### Rhabditidae UCE Analysis

Analysis of Ultra-Conserved Elements (UCEs) in Rhabditidae was performed to understand the genetic conservation and predictive features within this nematode family. UCEs were categorized into three groups: (1) UCEs shared across all analyzed species, (2) the most frequently observed UCEs, and (3) UCEs identified as important features by the XGBoost machine learning model. This analysis aims to highlight UCEs with potential biological significance by identifying overlaps between these groups.

**Table 16:** Summary of UCEs shared across Rhabditidae, grouped by their presence in Shared in all species (Shared), the most (Frequent), and the Feature selection in the XGBoost model (XGBoost) categories.

| UCE | Groups where it appears |
| --- | --- |
| uce.512 | Shared, Frequent |
| uce.655 | Shared, Frequent |
| uce.4011 | Shared, Frequent |
| uce.1892 | Shared, Frequent |
| uce.739 | Shared, XGBoost |
| uce.1739 | Shared, XGBoost |
| uce.16032 | Shared, XGBoost |
| uce.13654 | Frequent, XGBoost |
| uce.16977 | Frequent, XGBoost |
| uce.7408 | Frequent, XGBoost |

The UCEs shared between two groups offer valuable insights. For example, UCEs shared between “Shared by all species” and “Most Frequent” (uce.512, uce.655, uce.4011, uce.1892) indicate highly conserved and recurrent regions within Rhabditidae. UCEs shared between “Shared by all species” and XGBoost (uce.739, uce.1739, uce.16032) or “Most Frequent” and XGBoost (uce.13654, uce.16977, uce.7408) highlight regions important for species classification and differentiation. No UCEs are present in all three groups.

##### Panagrolaimidae UCE Analysis

Analysis of Ultra-Conserved Elements (UCEs) in Panagrolaimidae was performed to understand the genetic conservation and predictive features within this nematode family. The UCEs were categorized into three groups: (1) UCEs shared across all analyzed species, (2) the 100 most frequently observed UCEs, and (3) UCEs identified as important features by the XGBoost machine learning model. This analysis aims to highlight UCEs with potential biological significance by identifying overlaps between these groups.

The UCEs found in all three groups (uce.9910, uce.194, uce.8674, uce.459, uce.8786, uce.323) are particularly significant as they represent regions that are not only conserved across all species but also frequently observed and identified as important predictors by the XGBoost model. This suggests these UCEs could play crucial roles in the fundamental biology and evolution of Panagrolaimidae. The UCEs shared between two groups also offer valuable insights. For example, UCEs shared between XGBoost and Most Frequent may indicate regions that are highly informative for species classification, while those shared between XGBoost and Shared by all species might highlight regions under strong selective pressure.

**Table 17:** Summary of UCEs shared across Panagrolaimidae, grouped by their presence in Shared in all species (Shared), the most (Frequent), and the Feature selection in the XGBoost model (XGBoost) categories.

| UCE | Groups where it appears |
| --- | --- |
| uce.9910 | Shared, Frequent, XGBoost |
| uce.194 | Shared, Frequent, XGBoost |
| uce.8674 | Shared, Frequent, XGBoost |
| uce.459 | Shared, Frequent, XGBoost |
| uce.8786 | Shared, Frequent, XGBoost |
| uce.323 | Shared, Frequent, XGBoost |
| uce.10196 | Shared, XGBoost |
| uce.5837 | Shared, XGBoost |
| uce.6806 | Shared, XGBoost |
| uce.7309 | Shared, XGBoost |
| uce.9348 | Shared, XGBoost |
| uce.10095 | Shared, Frequent |
| uce.10105 | Shared, Frequent |
| uce.10397 | Shared, Frequent |
| uce.1316 | Shared, Frequent |
| uce.1795 | Shared, Frequent |
| uce.281 | Shared, Frequent |
| uce.4162 | Shared, Frequent |
| uce.431 | Shared, Frequent |
| uce.4329 | Shared, Frequent |
| uce.4462 | Shared, Frequent |
| uce.4491 | Shared, Frequent |
| uce.5306 | Shared, Frequent |
| uce.5355 | Shared, Frequent |
| uce.5498 | Shared, Frequent |
| uce.5515 | Shared, Frequent |
| uce.604 | Shared, Frequent |
| uce.618 | Shared, Frequent |
| uce.7157 | Shared, Frequent |
| uce.7646 | Shared, Frequent |
| uce.7753 | Shared, Frequent |
| uce.7765 | Shared, Frequent |
| uce.8373 | Shared, Frequent |
| uce.8702 | Shared, Frequent |
| uce.9422 | Shared, Frequent |
| uce.10015 | Frequent, XGBoost |
| uce.10136 | Frequent, XGBoost |
| uce.215 | Frequent, XGBoost |
| uce.4163 | Frequent, XGBoost |
| uce.4381 | Frequent, XGBoost |
| uce.451 | Frequent, XGBoost |
| uce.5260 | Frequent, XGBoost |
| uce.574 | Frequent, XGBoost |
| uce.9218 | Frequent, XGBoost |
| uce.9721 | Frequent, XGBoost |

## 1 Funding statement and acknowledgments

This research was conducted under the DFG funded project B08 in the CRC1211 [grant number 268236062] awarded to P. H. Schiffer and A-M Waldvogel, the ENP grant awarded to P. H. Schiffer [grant number: 434028868] and under the Biodiversity Genomics Center Cologne (BioC2) project funded by the UoC forum under the Excellent Research Support Program of the University of Cologne. We would like to thank the Cologne Center for Genomics (CCG) for their support, as well as Marie-Anne Félix for collecting and providing nematode cultures.

## 2 Conflict of interest statement

The authors declare that they have no conflict of interest.

## 3 Data availability statement

The sequencing data will be made available through the European Nucleotide Archive (ENA) and can currently be found in the Zenodo repository (10.5281/zenodo.15395838). The bioinformatic pipeline developed for machine learning genera classification is available on GitHub (https://github.com/LucyJimenez/uceml-analysis). Slides for the following nematodes used in the target capture analysis have been deposited at the Swedish Museum of Natural History: *Neocephalobus halophilus* (SMNH-226079–SMNH-226083); *Panagrolaimus* sp. PS1579 (SMNH-226085); *Panagrolaimus* sp. JU1371 (SMNH-226086); *Panagrolaimus* sp. JU1387 (SMNH-226087); *Panagrolaimus* sp. JU1645 (SMNH-226088); *Acrobeloides* cf. *guoghiensis* ARO.22.05 (SMNH-226113); *Acrobeloides tricornis* PAP.22.17 (SMNH-226112); *Panagrolaimus* sp. ALT.22.04 (SMNH-226089); *Panagrolaimus* sp. ALT.22.08 (SMNH-226090); *Panagrolaimus* sp. PAP.22.29 (SMNH-226091); *Panagrolaimus* sp. PAP.22.39 (SMNH-226092–SMNH-226083).

## Notes

### Competing Interest Statement

The authors have declared no competing interest.

### Summary of Updates

Typos have been fixed, tanglegram has been improved, and an extra phylogeny is included in the supplementary material. The models were re-run after noticing an error in the test data set, this improved the model results

